# Locating the Missing Chlorophylls *f* in Far-red Photosystem I

**DOI:** 10.1101/2024.08.06.606606

**Authors:** Giovanni Consoli, Fiazall Tufail, Ho Fong Leong, Stefania Viola, Geoffry A. Davis, Daniel Medranda, Nicholas Rew, Michael Hofer, Paul Simpson, Marco Sandrin, Benoit Chachuat, Jenny Nelson, Thomas Renger, James W. Murray, Andrea Fantuzzi, A. William Rutherford

**Affiliations:** Department of Life Sciences, Imperial College; London, SW7 2AZ, United Kingdom; Department of Physics, Imperial College; London, SW7 2AZ, United Kingdom; Institute for Theoretical Physics, Johannes Kepler University; Linz, 4040, Austria; Centre for Structural Biology, Imperial College; London, SW7 2AZ, United Kingdom; Sargent Centre for Process Systems Engineering, Department of Chemical Engineering, Imperial College London; London SW7 2AZ, United Kingdom

## Abstract

The discovery of chlorophyll *f*-containing oxygenic photosynthesis, with its long-wavelength photochemistry, represented a new low-energy paradigm. However, subsequent structural studies on chlorophyll *f*-containing Photosystem I (PSI) found five chlorophylls *f* but none among the photochemically active pigments and concluded that chlorophyll *f* plays no photochemical role. Here we report a cryo-EM structure (2.01 Å) of far-red PSI from *Chroococcidiopsis thermalis* PCC 7203, showing all eight chlorophylls *f*, including the redox active A_-1B_. Simulations of absorption difference spectra induced by charge separation indicate that the A_-1B_ chlorophyll *f* absorbs at 755 nm. The chlorophyll *f* sites, some wavelength assignments, and conserved far-red-specific amino acids, provide functional insights, including redox tuning of chlorophyll *f* as the primary donor and far-red excitation energy-sharing over the PSI trimer.

## Main Text

Chlorophyll *f* (Chl *f*) was discovered in a cyanobacterium isolated from stromatolites when grown under far-red light (*1*). The ability to produce Chl *f* is present in a small but diverse proportion of cyanobacterial species (*2, 3*) and is associated with a ∼21 gene cluster that codes for far-red (FR) light versions of the main chlorophyll-containing subunits of Photosystem I and II (FR-PSI and FR-PSII), as well as other proteins needed for FR-photosynthesis (*2*). In cells adapted to FR-light, Chl *a* still dominates, with only 7-8 Chl *f* molecules present in FR-PSI, and 4 Chl *f* and 1 Chl *d* in FR-PSII (*4*).

Originally, the long-wavelength pigments were thought to play only a light harvesting role (*5*), with heat from the environment needed for uphill excitation energy transfer to the higher energy, photochemically active, Chl *a* primary donor. This view was contradicted by biophysical studies of the FR-photosystems providing a range of evidence for long-wavelength primary electron donors in both FR-photosystems, most tellingly efficient FR-light-driven charge separation at cryogenic temperatures (*4*). Chl *f*-based photochemistry was a surprise and was greeted with excitement, as a new, low energy and potentially more energy-efficient paradigm for photosynthesis, one which resonated with the long-standing desire for more efficient crop production (*6*).

The primary donor in FR-PSII was proposed to be one of the 5 long-wavelength pigments and located in the Chl_D1_ position (*4*) and this was confirmed by cryo-EM to be the single Chl *d* (*7*). When cryo-EM structures of FR-PSI appeared, however, they showed no evidence for Chl *f* among the redox active pigments (*8*–*11*) and it was argued that Chl *f* had no photochemical role in this photosystem. Subsequent spectroscopic studies based their interpretations on this conclusion (*12*–*14*) and to reconcile the low-temperature, long-wavelength photochemistry with the absence of a Chl *f* in the reaction centre, an *ad hoc* model was proposed in which excitation energy is transferred from Chl *f* antenna pigments to a putative enhanced long-wavelength charge-transfer absorption band assigned to a Chl *a* pair in the reaction centre (*13*).

The only chemical difference between Chl *a* and Chl *f* is the substituent at the C2 position: a methyl group in Chl *a* and a formyl group in Chl *f* (*1*). In cryo-EM, the atomic Coulomb potential, which arises from the negatively charged electron cloud and the positively charged nucleus, scatters the electron beam resulting in a map of the electrostatic potential (ESP). Since the formyl oxygen of Chl *f* is negatively polarized, it contributes less to the signal (*15*–*17*). This property is reflected in previous cryo-EM models (*8*–*11*), the most recent of which classify 5 Chl *f* molecules as likely candidates from direct observation of additional ESP (*10*). Chl *f* assignments have also been influenced by consideration of the specific conserved changes in the FR-PSI proteins relative to their homologues in white light-absorbing Chl *a* PSI (WL-PSI), and by the chemistry of Chl *f*, i.e., the likely H-bond partners to its formyl group and its preference for a non-histidine axial ligand to the Mg^2+^(*4, 18*).

Given the intrinsic difficulties in detecting Chl *f* by cryo-EM, higher resolution structures are needed to solve the key mechanistic controversy over the presence of a Chl *f* among the redox active pigments of FR-PSI. To address this question, we have obtained a cryo-EM map of FR-PSI from *Chroococcidiopsis thermalis* PCC 7203 (*C. thermalis*) at 2.01 Å resolution and introduced an improved analysis pipeline to classify chlorophyll substituents (fig. S1). The electrostatic potential from the formyl group, FR-conserved amino acid sequence changes, and electrochromic shifts induced by P700^+^ on the nearby pigments, all provide evidence for the presence of Chl *f* in the A_-1B_ site, a candidate as the primary electron donor in PSI (*4, 19*–*21*).

### Location of the chlorophyll *f* sites in the cryo-EM structure of FR-PSI

The improved resolution allows the formyl groups to be detected from the ESP map for all eight Chl *f* molecules in *C. thermalis* FR-PSI: A_-1B_, A21, A23, B07, B30, B37, B38, and L03 (Fig.1 and 3, fig. S2-S7, supplementary text S1). The previous published structures allowed the assignment of some of the expected eight Chl *f* molecules, some of which are in common with the candidates presented here (*8, 9, 11*). The presence of a C2 formyl H-bond in five of these cases (A21, A23, B07, B30 and B37) likely contributes to the ESP signal by fixing the possible orientations of the formyl group and by diminishing the charge on the negatively polarised formyl oxygen atom. The assignments are supported by a new statistical analysis developed here that is based on the “cone-scan” method of Gisriel et al. (*16, 22*). The original cone-scan method analysed variations in ESP around the putative formyl carbon at the precise distance corresponding to the length of a carbon-oxygen double bond. However, the maximum variations in ESP (positive and negative) are not necessarily expected to occur at one fixed distance, therefore the approach used here quantifies the local ESP environment of chlorophyll substituents over a range of distances from their respective carbon atom, allowing a more thorough exploration of the ESP map and thus a more rigorous identification of the formyl groups (fig. S1).

**Fig. 1:**
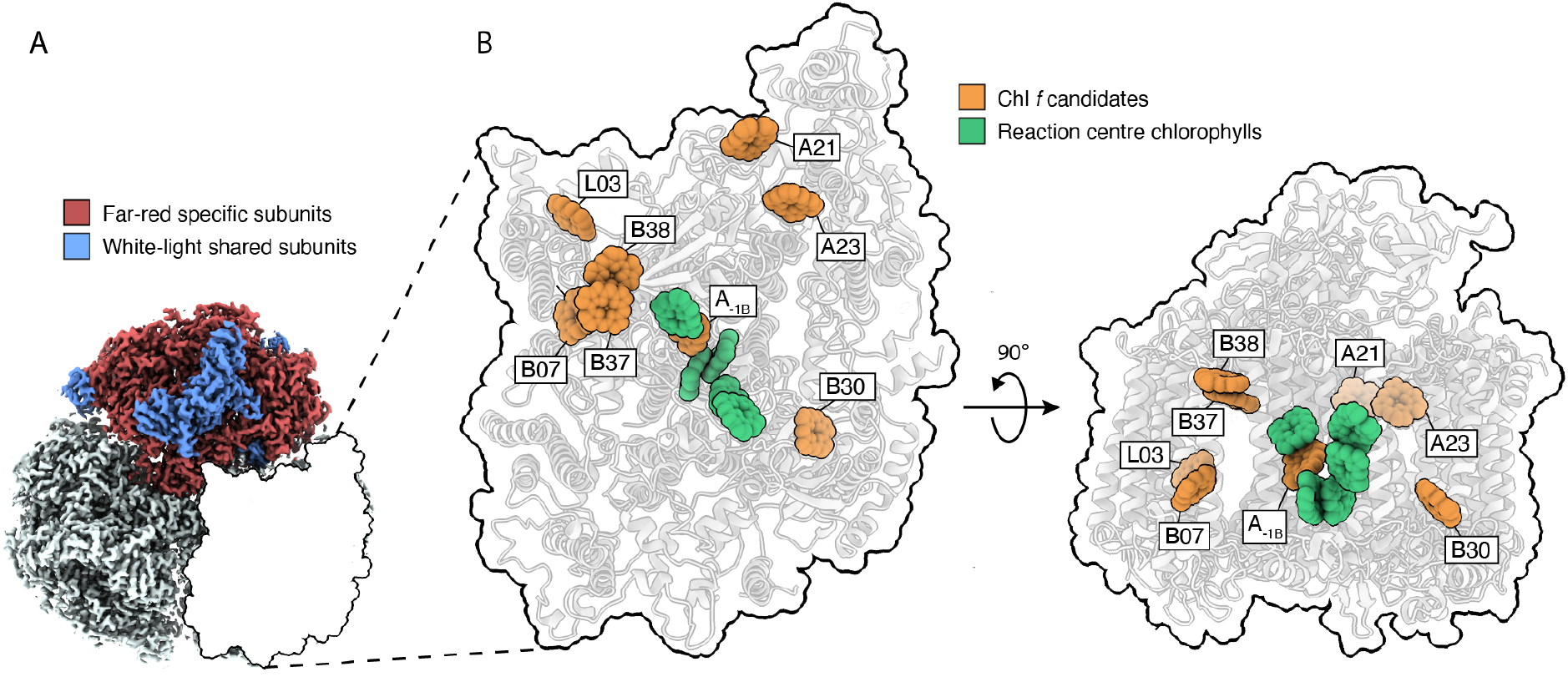
Structure of trimeric FR-PSI and chlorophyll *f* locations. **(A)** View of trimeric FR-PSI map from the cytoplasmic side. The map presents clear ESP for all the known PSI subunits: the six far-red light specific subunits (PsaA2, PsaB2, PsaF2, PsaI2, PsaJ2, PsaL2) are represented in red, while the six subunits in common with white light-PSI (PsaC, PsaD, PsaE, PsaK, PsaM, PsaX) are represented in blue.**(B)** Location of the 8 candidate Chl *f* molecules (in orange) in a FR-PSI monomer, as viewed from the cytosolic face and membrane plane. The 6 redox active chlorophylls are shown in the centre of the monomer (in green when Chl *a*), Chl *f* is in the A_-1B_ site while the 5 others are Chl *a*.

### Evidence for chlorophyll *f* in the A_-1B_ site in the reaction centre

The higher resolution cryo-EM data is used to map the ESP distribution over a conical surface around the C2’ carbon of the A_-1B_ chlorophyll. A positive ESP feature (red), evident at a torsion angle of ∼145° (see materials and methods), indicates the presence of a formyl group at the C2 position of the A_-1B_ chlorophyll and confirms the presence of Chl *f* among the 6 reaction centre chlorophylls (fig. S8), as originally suggested (*4*). A negative ESP feature blue/blue-green) is also present at ∼270°, at the interface between the formyl carbon-oxygen double bond and the benzyl ring of PsaA2 Phe^459^ (Fig. 2A, 2B, 2C, 2D). A survey of high-resolution structures of FR-PSI and WL-PSI in the literature shows that this negative feature is present in a similar location in other FR-PSI structures, i.e., *Halomicronema hongdechloris* (PDB: 6KMX) (*9*), *Synechococcus* sp. PCC 7335 (PDB: 7S3D) (*11*), but absent from the equivalent site in WL-PSI maps of comparable resolution (fig. S9).

**Fig. 2:**
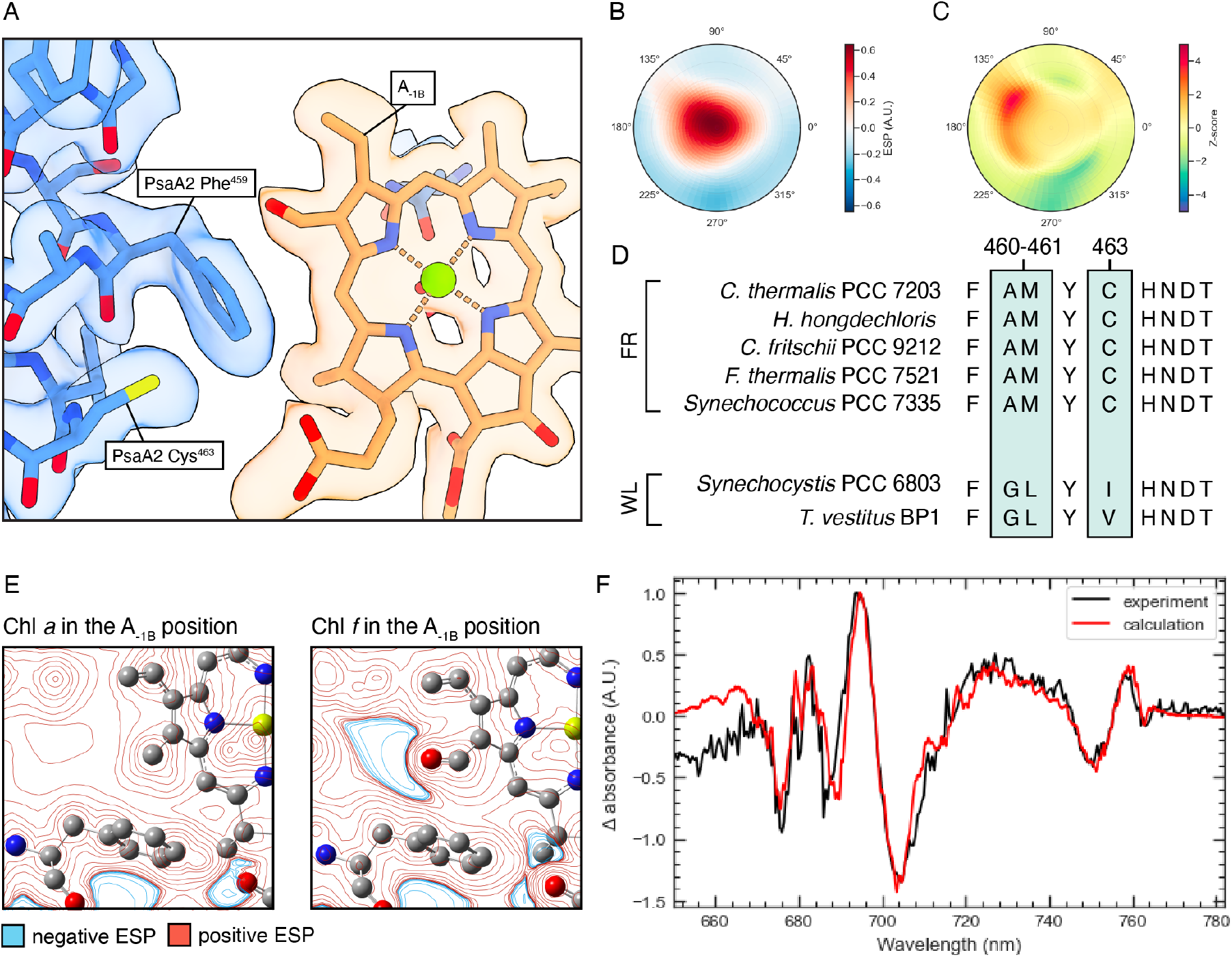
Evidence for A_-1B_ being a Chl *f*. (**A**) The chemical environment of the A_-1_ Chl on the B branch of the FR-PSI reaction centre. The Chl density is in orange, the density of the amino acids from PsaA2 in blue. (**B**) The ESP polar plot showing the presence of an increased positive potential at 145° torsion angle around the C2 substituent. (**C**) The Z-score polar plot highlighting the presence of a statistically significant (Z-score > 3) increase in positive (145° torsion angle) and negative (270° torsion angle) ESP features around the C2 of the A_-1B_ Chl relative to the averaged ESP environment around a methyl substituent (fig. S1). (**D**) The relevant part of the multiple sequence alignment of FR and WL PsaA, showing the far-red specific changes associated with the A_-1B_ site. (**E**) DFT calculated electrostatic environment around the C2 of A_-1B_, for Chl *a* (left) and Chl *f* (right) at the A_-1B_ site. Isopotential contour lines are shown on a plane defined by the A_-1B_ formyl group (C=O) and Phe^459^ γ-carbon, positive in red and negative in blue. Further details provided in fig. S10. (**F**) Comparison between the experimental (black) and the calculated (red) P700^+^ *minus* P700 difference spectra (at T=1.8 K).

Density Functional Theory (DFT) calculations of the electrostatic environment of A_-1B_ and its neighbouring amino acids are shown as isosurface contours of the ESP (Fig. 2E). A negative region (in blue) is present between the formyl group and the benzyl ring of PsaA2 Phe^459^ when Chl *f* occupies the A_-1B_ site. This negative region appears to correspond to the negative ESP feature detected at a polar angle of 270° in the ESP and Z-score polar plots (Fig. 2B, 2C) (see supplementary text S2).

The apparent lack of conserved amino acid changes near the A_-1_ sites in FR-PSI has been used as an argument against the presence of a Chl *f* at this location (*8*). However, a conserved change, from Ile/Leu/Val (depending on the species) to PsaA2 Cys^463^, is in fact present only in FR-PSI (Fig. 2D). The conserved sulfhydryl group of PsaA2 Cys^463^ and the C2 formyl of Chl *f* A_-1B_ are separated by the benzyl ring of PsaA2 Phe^459^ (Fig. 2A). The SH–π−formyl motif is likely to influence Chl *f* binding and redox tuning. The negative ESP near A_-1B_ (Fig. 2B, 2C, 2E), which is modified by the presence of the phenylalanine/cysteine pair (fig. S10), would be expected to destabilize the LUMO of the Chl *f* and make its redox potential more negative (*23*). This may compensate the intrinsically less reducing excited state of Chl *f* compared to that of Chl *a*, thereby restoring the reducing power needed for the PSI primary donor (*23*–*25*). Other conserved changes are present near the A_-1B_ site, PsaA2 Ala^460^ and PsaA2 Met^461^ with possible functional significance (Fig. 2D, fig. S11).

Fig. 2F shows a calculated P700^+^ *minus* P700 difference spectrum, in which the ground and excited states of the chlorophylls *a* and *f* were represented by atomic partial charges obtained from a fit of the electrostatic potential of their respective charge densities obtained with (time-dependent) density functional theory. An accurate description of the electrochromic shifts required i) the application of theory that goes beyond the point-dipole approximation, and ii) the specific physicochemical properties of Chl *a* and Chl *f* to be taken into account (material and methods).

The P700^+^-induced shifts on either Chl *a* or Chl *f* in the A_-1B_ site were found to be 17 cm^-1^ and -51 cm^-1^ respectively. This sign difference between the two types of chlorophyll was not previously predicted (*4, 26*). With Chl *f* in the A_-1B_ site and with a site energy at 747 nm (corresponding to an absorption at 755 nm), the improved calculation gave a P700^+^ *minus* P700 difference spectrum that almost quantitatively reproduced all features of the experimental spectrum (Fig. 2F). The difference spectrum could not be reproduced when the calculation was done with Chl *f* in any of the antenna chlorophyll locations (fig. S13, S14) or in the A_-1A_ site (fig. S15). An additional small but distinct negative peak around 762 nm was fitted as a band-shift from a Chl *f* antenna pigment. The largest electrochromic shifts of the Chl *f* pigments in the antenna arose from the B30 (7 cm^-1^) and B07 (10 cm^-1^) sites. Whereas B07 is excitonically coupled to L03, B30 is relatively isolated from the other Chl *f* pigments and its assignment to 762 nm would be consistent with the lack of a CD signal (*4*). Thus, the more likely candidate for the small negative dip at 762 nm in the P700^+^ *minus* P700 difference spectrum is B30. Support for the tentative assignment of B30 to the 762 nm absorption is obtained from the calculated difference spectrum (Fig. 2F), which reproduces the experimental spectrum (*4*).

The reports of a weak (*4*) or absent (*9*) Chl *f* bandshift in room-temperature difference spectra can be explained by the temperature dependence of the dielectric constant (fig. S16). Whereas the high-energy Chl *a* side of the difference spectrum is influenced both by the excitonic couplings (fig. S17) and the electrochromic shifts in site energies (fig. S18), the low energy Chl *f* region is dominated by electrochromic effects because Chl *f* at A_-1B_ is off-resonant to the other reaction centre pigments, which are Chl *a* molecules (fig. S19-S23) (see supplementary text S3).

### Consequences of chlorophyll *f* in the A_-1B_ site

Fig. 2 provides evidence that Chl *f* is in the A_-1B_ site, and it seems likely that it acts as the primary electron donor (*4*). Excitation with far-red light will result in charge separation mainly on the B-side of the reaction centre and this should be more marked at low temperatures. Most of the antenna Chl *f* molecules are located on the B-side, closer to A_-1B_, and this reflects the greater number of conserved amino acid differences in PsaB2 compared to PsaA2 (*4*). When excited by visible light, the Chl *a* antenna will mainly pass the excitation onto the Chl *f* pigments as seen in ultrafast kinetic studies (*12, 27*). Visible light may also excite redox active Chl *a* molecules (A_-1A_, P_A_ and P_B_), especially when they are excited directly, and these species are potentially capable of charge separation on the A-side (*4, 12, 27*) if primary electron transfer can compete with energy transfer to the low energy Chl f at A_-1B_. This predicted complex mixture of states may be unravelled by appropriate use the excitation wavelength selectivity.

No changes are seen in the vicinity of the A-side quinone, A_1A_, but a specific far-red conserved change around the lower potential quinone A_1B_ is present, the substitution of PsaB2 Ser^673^ with Thr (*28*). The structure around the quinone appears to be unaffected by this Ser to Thr change, thus a change in the redox properties of the A_1B_ is not expected. However, PsaB2 Thr^673^ is involved in a steric interaction with A_0B_ and this could limit any potential dynamic changes associated with quinone reduction. As pointed out recently, the PsaB2 Ser to Thr^673^ change, together with 2 other conserved amino acid changes (PsaB Val/Ile^666^ and Ser^667^), contribute to a 5° rotation of Chl A_0B_, changing its aromatic overlap with A_-1B_ and thus modifying their coupling (*28*). Quantum chemical calculations show that the rotation of Chl A_0B_ leads to increased electronic coupling in the Chl *f* A_-1B_/Chl *a* A_0B_ pair (see supplementary text S4 and fig. S24). This change in coupling may well be important for charge separation between Chl *f* A_-1B_ and Chl *a* A_0B_.

### Assignment of absorption peaks to chlorophyll *f* sites

A23 is assigned here as an H-bonded Chl *f*, in agreement with earlier work (*9, 10*) (fig. S3). The conserved amino acid changes in FR-PSI around this site in *C. thermalis* and *H. hongdechloris* are present in a sub-class of FR-PSI found in approximately half of FR-species, while the other half lack these changes and have a methionine instead of a glutamine in PsaA2 position 365 (fig. S3). In the structures of *F. thermalis* PCC 7521 and *Synechococcus sp*. PCC 7335, where PsaA2 Met^365^ is present, this chlorophyll was assigned as a Chl *a* based on sequence analysis and structure (*11*). A comparison of the 77 K absorption spectra in the literature from the 2 types of FR-PSI shows several differences in the 740-760 nm region (*4, 9, 13*) corresponding to changes in contributions of some Chl *f* molecules to the absorption spectra (i.e. excitonic coupling, reorganization energy) and the loss a Chl *f* in the A23 site. The complexity of the spectral differences does not allow a definitive interpretation, but it seems consistent with contributions from A23 to the absorption around 750 nm and an underlying absorption from A_-1B_ Chl *f* at 756 nm (see supplementary text S1).

77 K fluorescence spectra of FR-PSI show the presence of two terminal emitters, one at ∼750 nm and the other at ∼800 nm, the exact wavelength depending on the species (*4, 13*). The two terminal emitters are expected to be separated from each other, both spatially and by an energy barrier, such as by several Chl *a* molecules. Tros et al. (*29*) showed that while the emitter at ∼750 nm was insensitive to the presence of P700^+^, the emitter at ∼800 nm was quenched by P700^+^. The emission at ∼800 nm originates from a coupled pair of Chl *f* molecules, B37 and B38 (*4, 13*) with a distance from P^B+^ of 26 Å. In this pair, the inter-pigment distance is below 4 Å, where short-range contributions to site energy shifts are expected to be significant (*30*). The quenching of the ∼800 nm emitter by P700^+^ seems likely to be protective in excess light.

The lack of quenching of the ∼750 nm emitter likely reflects a larger distance from P700^+^ than the 800 nm emitter. From their locations (fig. S23), A21 and B30 are the most likely candidates as the emitter at ∼750 nm, as already suggested (*13*), as they are both far from P700^+^ (34 Å and 28 Å) and seem relatively isolated from the central core of Chl *f* molecules. However, the wavelength assigned to B30 is 762 nm based on its contribution to the band-shifts in the difference spectrum (Fig. 2F). Therefore, A21 appears to be the more likely of the two candidates. Given the proximity of A21 to A23, for A21 to be the terminal emitter, A23 would have to absorb at a shorter wavelength within the range assigned to it above (740-760 nm), otherwise A21 would transfer excitation energy to the lower energy pigment (A23) rather than emit.

While individual Chl *f* molecules can be tentatively assigned to specific absorptions in the spectrum (A_-1B_, 755 nm, B37/B38 pair at 800 nm, B30 at 762 nm), there is insufficient evidence to assign the others. Now they have all been localized, future work will focus on their physical and functional properties.

### Functional significance of the trimerization on excitation energy transfer

ESP polar plots of Chl L03 shows a clear density for a formyl group in the C2 position that does not appear to be H-bonded. A FR-specific conserved amino acid change is present, with PsaL2 Val^81^ replacing the Lys/Leu/Ile of the white light PsaL, generating the space necessary for the formyl group. The increased ESP density is present at a polar angle that suggests a possible interaction with a diglyceride present both in WL and FR-PSI (Fig. 3A, 3B, 3C).

**Figure 3:**
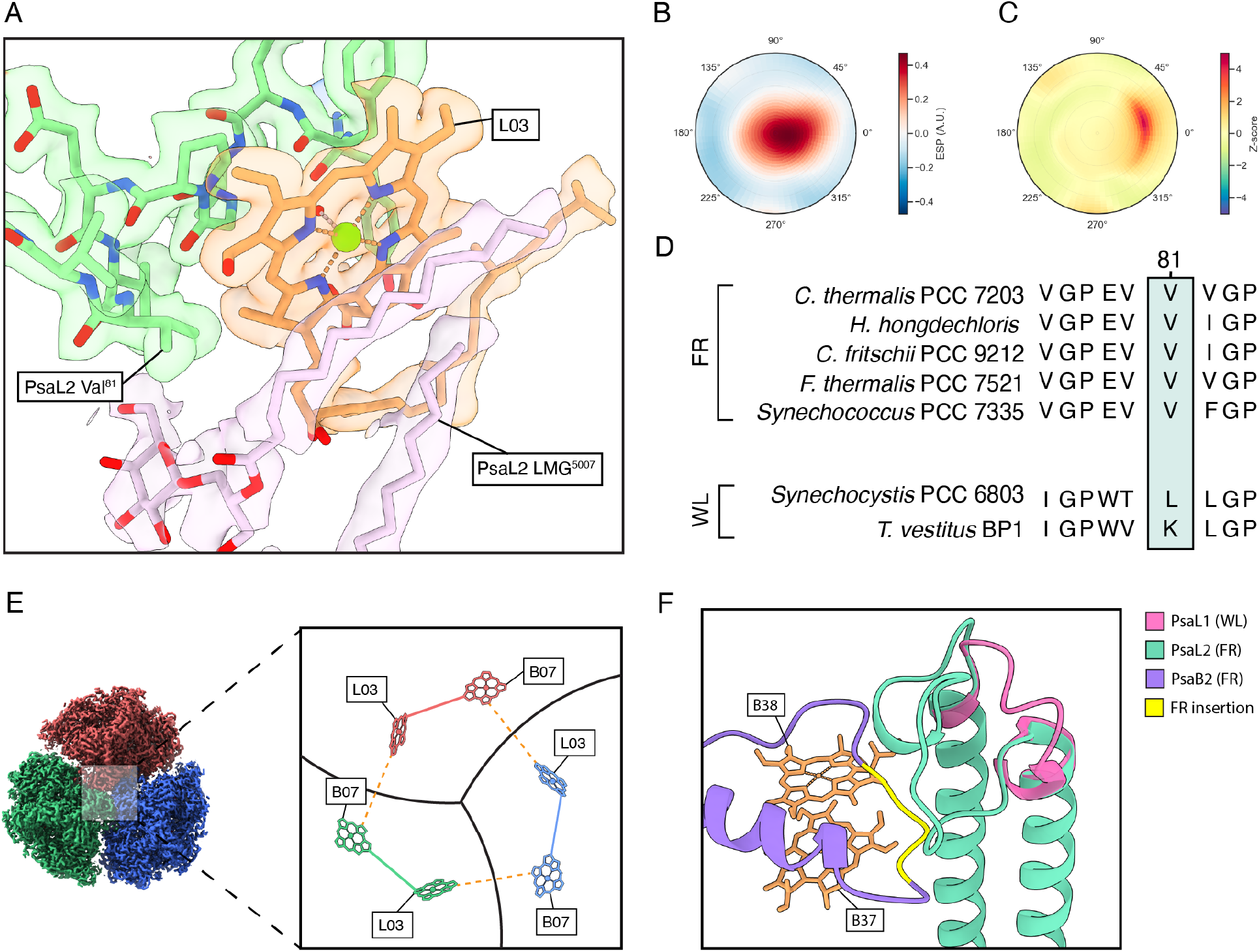
L03 Chl *f* site, a long wavelength bridge between the monomers. (**A**) Chemical environment of the L03 Chl. The Chl density is in orange, the density of the amino acid residues from PsaL2 in green, the density of the diglyceride in pink, and the density from water in red. (**B**) ESP polar plot indicating the presence of an increased potential around the C2 substituent. (**C**) Z-score polar plot indicating the presence of an increased potential around the C2 substituent. (**D**) Multiple sequence alignment of FR and WL PsaL, showing the far-red specific Val residue in position 81. (**E**) Lumenal view of trimeric FR-PSI, each monomer of the trimer coloured in red, green and blue, and zoom on the Chl *f* sites assigned near the monomer interfaces (B07 and L03). The intra-monomeric distance (solid lines) between the two pigments is ∼11 Å and the inter-monomeric distance (dashed lines) is ∼15 Å. (**F**) Structure of the FR-PSI specific PsaB2 insertion (yellow) near the B37/B38 Chl *f* dimer.

FR-PSI exists as trimers *in vivo*, whereas WL-PSI is found as dimers or tetramers in *C. thermalis* (*31*). The Chl *f* sites identified here are mainly located on the PsaB2 side of the FR-PSI complex and trimerization results in that side being close to the interface between individual PSI monomers. Thus, the formation of FR-PSI trimer results in a core structure enriched in Chl *f* (fig. S25). PsaL2 mediates trimer formation and the Chl *f* in the L03 site is between the B07 Chl *f* of the same complex and that of the neighbouring one. Thus, the Chl *f* L03 likely plays an important role in the Chl *f* connectivity in FR-PSI trimers. Within this core, the distances between Chl *f* molecules could allow efficient excitation energy transfer between the Chl *f* molecules and between the PSI monomers in the trimer (Fig. 3E). Excitation energy sharing between the individual PSI complexes in the PSI trimer has been suggested to occur in WL-PSI via specific routes, one of which involves the chlorophylls of PsaL (*32*). Excitation energy sharing among the Chl *f* molecules within the trimer would increase the chances of charge separation when one or both other PSI monomers in the trimer are photochemically “closed”.

The importance of this inter-trimer Chl *f* connectivity is also highlighted by another structural feature: the FR-specific B37/B38 loop of PsaB2, which has extensive interactions with a FR-specific loop of PsaL2 (Fig. 3F). These interactions could stabilize the Chl *f* dimer, B37/B38, favour the association of PsaB2-PsaL2, and prevent the formation of a chimeric WL-FR-PSI by obstructing PsaL1 and PsaB2 interactions.

## Conclusions

The structural model here shows the positions of all the expected Chl *f* sites in FR-PSI. H-bonds stabilise most of the antenna Chl *f* molecules, i.e., A21, A23, B07, B30, B37 (fig. S2-S6), making the formyl oxygens on Chl *f* more easily detectable by cryo-EM. When the formyl oxygens are not H-bonded, i.e., A_-1B_, B38 and L03 (Fig. 2, 3, and S7), they are more difficult to detect because of their increased negative partial charge and because they may occupy a broader range of torsion angles. This class of formyl oxygens therefore required improved resolution and the development of more thorough statistical analysis to allow them to be detected in cryo-EM maps. This explains their apparent absence in earlier work.

The present work also confirms that both positive and negative ESP signals can provide information on the electrostatic environment and the orientation of functional groups of cofactors and amino acid sidechains, as discussed previously (*15*–*17*). Here information obtained from this extra data provides additional mechanistic insights. When present, the H-bonds to the formyl oxygen not only stabilise the binding of the Chl *f*, but also shift the redox properties of the chlorophylls, making them easier to reduce and harder to oxidise. This would not greatly affect the antenna function of chlorophylls (other than small changes in their absorption peaks), but it would be detrimental for Chl *f* when it comes to a redox role, and particularly as the primary donor, which is the most reducing species in PSI, the low potential photosystem. Therefore, a Chl *f* acting as a primary donor in PSI would be expected to lack an H-bond to its formyl group and thus be difficult to detect by cryo-EM. Both expectations were found to be the case for the Chl *f* in the A_-1B_ site. In fact, the FR-A_-1B_ appears to be specifically negatively charged (Fig. 2B, 2C, 2E), an indication of redox tuning.

The identification of Chl *f* at the L03 site in the core of the trimer and at the A_-1B_ site in the FR-PSI reaction centre, represent significant advances in understanding how FR-PSI works. The L03 Chl *f* indicates exciton energy sharing between the PSI monomers in the trimer, while the A_-1B_ Chl *f* indicates photochemistry with Chl *f* as the primary donor. The previous spectroscopic studies of FR-PSI, which assumed no Chl *f* in the reaction centre, and their resulting mechanistic models of excitation energy transfer and charge separation, can now be re-assessed. FR-PSI is comparable to Chl *d* PSI in *Acaryochloris marina*, in that they both use near-IR photons (P740) for oxygenic photosynthesis and thus they have broken the red photochemical energy limit. As seen with PSII (*4, 33*), these two distinct low energy paradigms have evolved different strategies to do the same chemistry as in conventional Chl *a* photosynthesis but by using less energy. Comparisons of the two long-wavelength paradigms with each other and with Chl *a* photosystems, will advance our understanding of the mechanisms of all three. This knowledge could bring more efficient, long-wavelength crops a little closer.

## Acknowledgments

We thank Kenta Renard for the numerous discussions, Jan Schuller for the insightful conversation and suggestion on grid preparation and data processing, Doryen Bubeck for the useful suggestions on data processing, Ville Kaila for helpful suggestions on the computational approaches, and Suhail Islam for computation assistance, and Dominik Lindorfer and Sebastian Egginger for support in the early stage of the calculation of optical difference spectra. We thank the centre for structural biology at Imperial College for the training provided, for help with the early stages of sample screening and with data collection. We thank Diamond for the access and support of the cryo-EM facilities at the UK national electron Bio-Imaging Centre (eBIC), proposal BI25127.

## Funding

Marie Skłodowska-Curie grant agreement, No. 955520 – GC, AWR, AF, BC

Biotechnology and Biological Sciences Research Council, BB/R001383/1 – AWR, AF, SV, GAD

Biotechnology and Biological Sciences Research Council, BB/V002015/1 – AWR, AF, JWM, SV, DM

Biotechnology and Biological Sciences Research Council, BB/R00921X – AWR, AF, GAD

Biotechnology and Biological Sciences Research Council, BB/M011178/1 – FT

Leverhulme Trust Grant RPG-2022-203 – AWR, GAD, AF

European Research Council, Advanced Grant CAPaCITy, grant agreement no. 742708 – JN, DM

The Royal Society (Royal Society Research Professorship 2021) – JN, NR

Engineering and Physical Sciences Research Council Doctoral training award - NR

The Royal Society (Royal Society Research Professorship 2024) - AWR

Austrian Science Fund (FWF) P33155-NBL - TR

Austrian Science Fund (FWF) I-6313-N – TR

## Author contributions

GC, FT and HFL initiated the study.

SV grew and harvested *C. thermalis* PCC 7203

GC, FT, HFL, AF, and AWR conceived of the main experiments, collated results and interpretations, and wrote the article with input, edits, and approval from all authors.

SV and GAD developed and performed the isolation of the photosystems.

GC and FT prepared the grids.

GC, FT and PS collected the micrographs.

GC and FT processed the data to obtain the map. GC, FT and JWM built the atomic model.

HFL and AF collected the sequences and performed the phylogenetic analysis. GC and JWM developed and coded the local ESP quantification method.

DM, NR and JN performed DFT calculations.

MH and TR calculated the electrochromic shifts, simulated the P700+ *minus* P700 difference spectrum, and performed the underlying electronic structure calculations.

GC and BC fitted the low temperature absorption spectra.

## Competing interests

Authors declare that they have no competing interests.

## Data and materials availability

The atomic coordinates have been deposited in the PDB with accession code 9EYS. Data and scripts used in data processing, analysis, and visualization are available on GitHub (https://github.com/giovanniconsoli/Chlorophyll-subsituents-scan).

## Supplementary Materials

Materials and Methods

Supplementary Text S1: Chl *f* site assignments in common with literature

Supplementary Text S2: DFT calculation of the A_-1B_ chemical environment

Supplementary Text S3: Electrochromic shifts data and discussion

Supplementary Text S4: A_-1B_ /A_0B_ Electronic Coupling

figs. S1 to S25

tables S1 to S20

References (34-66)

## Notes

### Competing Interest Statement

The authors have declared no competing interest.

