## Supplementary Materials for "Locating the Missing Chlorophylls *f* in Far-red Photosystem I"

**The PDF file includes:**

Materials and Methods
Supplementary Text S1: Chl *f* site assignments in common with literature
Supplementary Text S2: DFT calculation of the $A_{-1B}$ chemical environment
Supplementary Text S3: Electrochromic shifts data and discussion
Supplementary Text S4: $A_{-1B}$ /$A_{0B}$ Electronic Coupling

Figs. S1 to S25
Table S1 to S20
References (34 - 66)

### Materials and Methods

Culture growth under far-red light of *C. thermalis* PCC 7203

*Chroococcidiopsis thermalis* PCC 7203 was grown in liquid BG11 medium (*34*) at 30 °C under 750 nm LED illumination at an intensity of ~30 μmol photons $m^{-2}$ $s^{-1}$ (750 nm, Epitex; L750-01AU). Cultures were harvested, pelleted and flash frozen after at least 3 months of exposure to exclusively far-red illumination.

Isolation of far-red-PSI complexes

Cells were broken with two passages in a flow cell disruptor (Constant Systems) at a pressure of 39 kPsi and then centrifuged at low speed (1000 g, 5 min at 4°C) to remove any unbroken material. Isolation was carried out in darkness or minimal dim green light. The supernatant was then centrifuged at 40,000 rpm in a Ti45 rotor at 4ºC for 20 minutes to pellet the membranes. Membranes were resuspended to 1 mg $mL^{-1}$ total chlorophyll in TM buffer (50 mM MES – NaOH pH 6.5, 5 mM $CaCl_2$, 10 mM $MgCl_2$, 1.2 M Betaine) and solubilized at 4º C for 1 hour by the addition of β-DM to a final concentration of 0.4% (w/v) in the dark. After the removal of insoluble debris by centrifugation, the supernatant was then concentrated (100 kDa Filters, Amicon) and the solubilized membranes were loaded onto sucrose gradients (50 mM MES – NaOH pH 6.5, 5 mM $CaCl_2$, 10 mM $MgCl_2$, 500 mM Sucrose, 0.04% β-DM) and centrifuged overnight at 120,000 x g at 4º C in a SW28 rotor with maximum acceleration and deceleration. Bands containing Photosystem I (PSI) trimers were harvested from the sucrose gradients and washed multiple times with TM buffer before being loaded on a MonoQ 5/50 anion exchange column (GE Healthcare). The column was washed with 5 column volumes of Buffer A (20 mM MES – NaOH pH 6.5, 5 mM $CaCl_2$, 5 mM $MgCl_2$, 0.03% β-DM). Fractions were eluted with a 0% to 100% gradient with Buffer B (Buffer A + 600 mM NaCl) at a flow rate of 0.5 ml/min over 50 minutes. Eluted peak fractions were pooled, washed to remove NaCl, concentrated (100 kDa Filters, Amicon) and stored at 4°C or flash frozen and stored at -80°C depending on the use.

Grid preparation

C-Flat 1 µm hole size and 1 µm hole spacing with a 300-copper mesh grids (Agar Scientific) were glow-discharged for 30 s at 25 mA and 3.5 µl of far-red (FR)-PSI sample (1 mg/ml of chlorophyll, equivalent to ~4.2 mg/ml of protein) were applied. Grids were blotted at 4°C and 100% humidity in the presence of only dim green light and plunge frozen in liquid ethane after 3 seconds with a Vitrobot mark IV (Thermo Fisher Scientific).

Data acquisition

Initial screening of the sample was conducted on a Glacios (Thermo Fisher Scientific) operated at 200k and a magnification of 75k. Particles were imaged using a Krios III (Thermo Fisher Scientific) operated at 300 kV and a magnification of 130k. Images were recorded on a Falcon 4i (Thermo Fisher Scientific) with a pixel size of 0.921 Å and a dose of 40 electrons per $Å^2$ for a total of 40 frames. Images were collected in super resolution mode with a SelectrisX energy filter with a slit width of 20 eV. The targeted defocus range was varied from –0.5 to –3.5 µm using the EPU software (Thermo Fisher).

Single particle analysis

A total of 8854 movies were collected from two different grids. The frames were aligned, dose weighted, and the contrast transfer function (CTF) was estimated in CryoSPARC v4.4.1 (*35*). Micrographs were curated by removing those with CTF fits below 10 Å. The subset obtained contained 7455 micrographs (84.1%) and was used to hand pick ~2000 PSI-type particles across the defocus range. Particles were 2D classified and used to template pick across the entire dataset, yielding 1,560,287 picks. After multiple rounds of 2D classification and *ab initio* refinement, duplicated particles were removed and a subset of 138,048 particles was used for homogeneous refinement to obtain an initial map >5Å and to confirm C3 symmetry. After multiple rounds of per particle CTF refinement and local motion correction, a set of 83,225 particles were used to perform non-uniform refinement imposing C3 symmetry, producing a map at a global resolution of 2.01Å based on GC-Fourier Shell Correlation at a cut-off of 0.142.

Model Building

The cryo-EM structures of a white light PSI (PDB 7FIX) (*36*)and a FR PSI (PDB 7LX0) (Gisriel et al., 2020) were fitted to the ESP map using the Phenix software suite (*37*). Rebuilding of the initial model was done with Coot (*38*) and then refined in real space in phenix.real_space_refine (*39*) and validated with MolProbity (*40*) (Table S1). Chl and carotenoid numbering are based on those reported in Jordan *et al.* (*41*).

Quantification and statistical analysis of the electrostatic potential

The electrostatic potential (ESP) is interpolated on the surface of a cone with its axis extending from that of the CX – CX' bond, where X is the IUPAC number given to the chlorin carbons that have relevant substitutions, and an aperture 120° ($2\theta$). The ESP is sampled within the range of distances 0Å to 2.5Å ($d$), with a step size of 0.1Å. This is repeated for every 5° rotation of the torsion angle ($\varphi$). The 90° torsion angle is defined as being parallel to the plane defined by the positions of the three carbon atoms, CX-1, CX and CX', in the direction of CX-1, where CX is the IUPAC carbon number on the chlorin ring. The direction of rotation of the torsion angle is clockwise by IUPAC convention.

Our implementation is written in Python using the GEMMI library (*42*). The Jupyter notebook computes the average ($\mu$) and standard deviation ($\sigma$) of the ESP on the surface of the cone for each of the substituents (C2, C3, C7, C8, C12). It thus provides visual information on the quality of the map and on the preferential orientation of the vinyl (C3) and ethyl (C8) substituents with respect to the chlorophyll plane. It also calculates the precision, recall and accuracy as metrics of the quality of the analysis. To classify the substituents in the C2 position, the Z-score of each sampling point is computed with respect of the average methyl substituent in position C7.

$$Z_{[d,\varphi]} = \frac{ESP^{C2}_{[d,\varphi]} - \mu^{C7}_{[d,\varphi]}}{\sigma^{C7}_{[d,\varphi]}}$$

The ESP and the Z-scores on the surface of the cones are plotted as 2D projection looking down the CX – CX' axis (fig. S1). This approach allows for the possibility that maximum differences between substituents could occur outside of the predicted length of the formyl's carbon – oxygen double bond. It thus explores a larger area compared with the original cone-scan method which sampled a single distance and gave a single line around the cone (*16*). Moreover, using Z-scores instead of the strict $\mu + 3\sigma$ cutoff as used previously (*10*, *16*). allows a more quantitative investigation of both positive and negative features in the environment of the substituent.
The code is available on GitHub at the following link (https://github.com/giovanniconsoli/Chlorophyll-subsituents-scan).

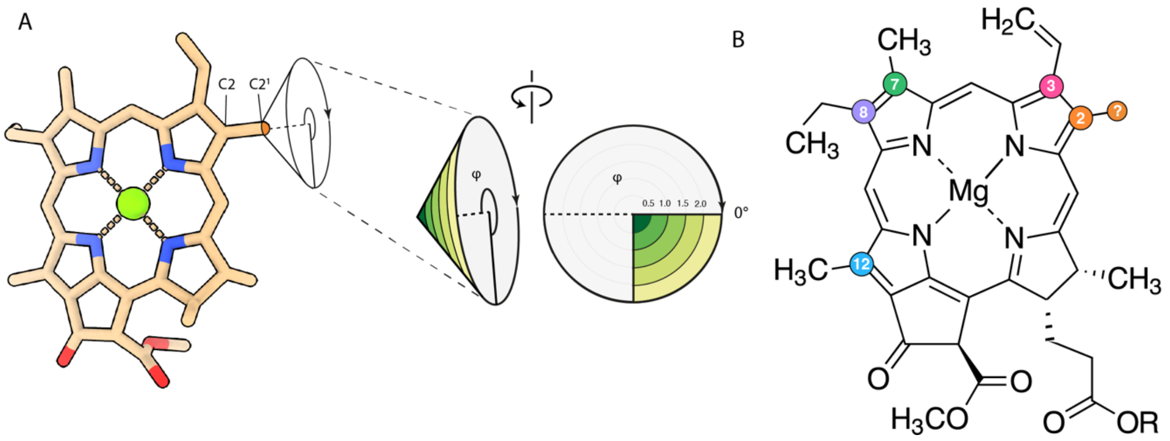


**fig. S1. Quantification and statistical analysis of the electrostatic potential**
(A) An example of geometric construction of the cone for substituent in the C2 position. The aperture is 120° and the range of distance scanned i 0-2.5 Å from C2'. (B) Location of the substituents used for the analysis are shown on the chemical structure of the chlorophyll. The C7 (green) methyl group is used to calculate the average methyl polar plot and its standard deviation, and these are used to calculate the Z-scores of the substituents (C2, C3, C8 and C12). The C3 (red) vinyl group and the C8 (violet) ethyl group are used as controls, and they should be differentiated by this method. The C12 (cyan) methyl is used as a negative control, i.e., it is expected to be indistinguishable from the C7 methyl group. The C2 (orange) methyl or formyl are the targets of the analysis. Any substituent with a Z-score above 3 is considered a positive result, i.e., significantly different from a methyl group.

Density Functional Theory

Single-point energy calculations were performed for eight different molecular assemblies derived from cryo-EM data for FR-PSI of *C. thermalis*. The assemblies were obtained using either, 1) structures fitted to the 2.01 Å resolution cryo-EM model; or 2), as per 1) but using structures in which the molecular components were replaced with those geometry optimised with DFT (details below). The structure fitted to the cryo-EM model shows a Chl *f* in the $A_{-1B}$ position, and the remaining chlorophyll sites distributed and identified as Chl *a* or Chl *f* as per Fig. 1. For case 1), cryo-EM data were post-processed as specified in Model Building. For case 2), each molecular component was embedded in a solvent of dielectric constant eps=2 with the Solvation Model based on Density (SMD) and geometry optimised individually using the B3LYP/6-31+G(d) level of theory. Post-optimisation, the optimised components were realigned to the 2.01 Å experimental structure in PyMol. In each case, the resulting alignment resulted in a RMSD between mobile and target objects less than 0.32 Å.

DFT calculations were performed for the $A_{-1B}$ – $A_{0B}$ chlorophyll pair located in the reaction centre ($A_{-1B}$ as the putative Chl *f*.) For the $A_{-1B}$ – $A_{0B}$ structures obtained using methods 1) and 2) as specified above, pairs were generated by duplicating the structure but with the C2 formyl group of Chl *f* in the $A_{-1B}$ position substituted by a methyl group *(i.e.*, converting it to Chl *a*), leading to four assemblies. A further four assemblies were generated by duplicating these four structures but with the PsaA $Phe^{459}$ and $Cys^{463}$ removed, taking the total number of assemblies to eight. This was done to identify the ESP signal expected from either Chl *a* or Chl *f* at that site, and the effect of the $Phe^{459}$ - $Cys^{463}$ motif on the local ESP environment.

For each assembly, the following were included: the chlorophyll of interest with phytol chain truncated up to the C1 carbon; the coordinating ligand to the $Mg^{2+}$ ion; and at least any amino acid residues within 3 Å of the C2 carbon. All amino acids were included up to the peptide bond. The remaining terminal carbons were methylated to ensure a singlet ground state for DFT calculations. The single point calculations were performed using the same level of theory as the optimisations, *i.e.,* B3LYP/6-31+G(d) with the molecular assemblies embedded in an eps=2 environment using the SMD model. Electrostatic potential (ESP) maps were generated using the cubegen utility in Gaussian. Cone scans of ESP were extracted from the computational data using an in-house script for comparison with the experimentally derived cone scans. All DFT calculations were carried out using Gaussian 16vC.01.

Charge Density Maps

Cryo-EM scattering factors are obtained by taking the Fourier transform of the ESP. However, cryo-EM maps are known to contain artefacts that appear as a function of scattering angle, ionisation state, and proton number (*17*). To aid interpretation of the experimental data, the cryo-EM ESP cone scans were additionally compared with cone scans of the electron density (ED) and charge density distributions (CD) from the computational data set. CDs are not obscured by the long-range effects apparent in ESP maps, and CD iso-surfaces allow for improved identification of sidechain rotamers since the maps more closely resemble the sum of images of independent atoms (*43*). CD distributions were obtained for each of the twelve computationally obtained ESP maps and the experimental ESP map in ChimeraX following the method of Wang (*43*).

**Calculation of optical difference spectra**

The P700$^+$ *minus* P700 absorption difference spectrum is defined as

$$\Delta\alpha(\omega) = \alpha(\omega)_{\mathrm{P+}} - \alpha(\omega)_{\mathrm{P}} \qquad (1)$$

where $\alpha(\omega)_{P^+}$ is the absorption spectrum of the complex with the positively charged (oxidized) chlorophyll pair (P700, made up of $P_A$ and $P_B$), and $\alpha(\omega)_P$ is the absorption spectrum with the neutral chlorophyll pair. The oxidized pair was created in the experiment by light excitation at 77K and the positive charge is assumed to be localized on pigment $P_B$ (*44–47*) Note, that we neglect the influence of the reduced iron-sulfur cluster, since it was shown to be small (*48*).

The exciton Hamiltonian relevant for the calculation of the spectra reads:

$$H_{\mathrm{exc}} = \sum_m E_m \, |m\rangle\langle m| + \sum_{m,n} V_{mn} |m\rangle\langle n| \qquad (2),$$

where $E_m$ is the site energy of pigment $m$ and $V_{mn}$ is the excitonic coupling between $m$ and $n$. The excitonic coupling $V_{mn}$ is calculated using the TrEsp-method (*30*, *49*) giving:

$$V_{mn} = f \sum_{I,J} \frac{q_I^{(m)}(1{,}0) q_J^{(n)}(1{,}0)}{|\boldsymbol{R}_I^{(m)} - \boldsymbol{R}_J^{(n)}|} + \delta_{m,\mathrm{P_A}} \delta_{n,\mathrm{P_B}} V_{mn}^{(CT)} \quad (3),$$

where the transition charge $q_I^{(m)}(1{,}0)$, which is placed on the *I*th atom of pigment *m* is obtained from a fit of the ESP of the transition density of this pigment, and *f* is a screening constant. The transition charges of Chl *a* obtained in this way where rescaled by a constant factor to reproduce the magnitude of the experimental transition dipole moment of 5.47 D measured in a solvent with optical dielectric constant $\varepsilon_{\mathrm{opt}} = 2$ (*50*), which represents the average optical dielectric constant of PSI (*51*). In this way we take into account implicit solvent effects on the transition dipole moment (*52*). For Chl *f*, we use the same correction factor as for Chl *a*, since no experimental investigation of its oscillator strength in different solvent environments is available yet. The average screening factor $f = 0.69$ was obtained from Poisson-TrEsp calculations on PSI (*53*). For the excitonic coupling within the P700 pair, we take into account a superexchange type contribution $V_{\mathrm{P_A P_B}}^{(\mathrm{CT})}$ arising from the coupling between local excited and charge transfer (CT) states, in addition to the long-range Coulomb coupling, resulting in an overall coupling of $V_{\mathrm{P_A P_B}} = 225$ cm$^{-1}$ (*30*). Please note that 84 % of this coupling is due to short-range contributions, due to wavefunction overlap (*30*). There are also a few closely spaced chlorophylls in the antenna (*54*), for which, however, there are no estimates of short-range effects available. Since these chlorophylls have no critical influence on the optical difference spectra calculated here, we neglect the short-range contributions to the excitonic couplings involving these antenna pigments (known as red chlorophylls). Note that there is one Chl *f* pair (B37/B38), which most likely gives rise to the redmost excited state at around 800 nm, as discussed in the main text. For this dimer we have taken into account the short-range effects in an effective way, by shifting the site energies of B37 and B38 to 800 nm. The details of the quantum chemical calculations of transition charges will be presented further below.

The site energies $E_m$ of the RC pigments and of the antenna pigments in close vicinity to the RC are determined from a fit of the experimental difference spectrum $\Delta\alpha(\omega)$, whereas the site energies of the remaining antenna pigments were chosen such that the overall inhomogeneous absorption spectrum agrees with experimental data. Explicit values for the excitonic couplings and site energies are given below. There is, of course, a large ambiguity concerning the site energies of many Chl *a* pigments in the antenna. However, these site energies have only a minor and indirect influence on the P700$^+$ *minus* P700 difference spectrum, used here to prove that there is Chl *f* bound to the $A_{-1B}$ site in the RC.

For the oxidized special pair state ($P^+$) the site energies of the remaining pigments in PSI

$$E_m(P^+) = E_m(\mathrm{P}) + \Delta E_m \qquad (4)$$

experience electrochromic shifts $\Delta E_m$ with respect to $E_m(\mathrm{P})$ for the neutral state (P). Using the charge density coupling (CDC) method (*55*) the electrochromic shift is obtained as:

$$\Delta E_m = \frac{1}{4\pi\epsilon_0\epsilon_{\mathrm{eff}}}\left(\left(V_{\mathrm{eg}_+}^{(m,\mathrm{P_B})} - V_{\mathrm{gg}_+}^{(m,\mathrm{P_B})}\right) - \left(V_{\mathrm{eg}}^{(m,\mathrm{P_B})} - V_{\mathrm{gg}}^{(m,\mathrm{P_B})}\right)\right)$$
$$= \frac{1}{4\pi\epsilon_0\epsilon_{\mathrm{eff}}}\sum_{I,J}\frac{\Delta q_I^{(m)}(\mathrm{e,g})\,\Delta q_J^{(\mathrm{P_B})}(\mathrm{g_+,g})}{|\boldsymbol{R}_I - \boldsymbol{R}_J|} + \delta_{m,\mathrm{P_A}}\Delta E_m^{(\mathrm{CT})} \qquad (5)$$

The CDC between the excited (ground) state of pigment *m* and the ground state of $P_B$, denoted as $V_{eg}^{(m,\mathrm{P_B})}$ ($V_{gg}^{(m,\mathrm{P_B})}$). These CDCs are expressed in terms of atomic difference charges, where $\Delta q_I^{(m)}(e,g) = q_I^{(m)}(e) - q_I^{(m)}(g)$ is the difference of the atomic partial charges between the excited and the ground state obtained from a fit of the electrostatic potential (ESP) of the charge densities of the two states. $\Delta q_I^{(\mathrm{P_B})}(g_+,g) = q_I^{(\mathrm{P_B})}(g_+) - q_I^{(\mathrm{P_B})}(g)$ is the difference of the partial charges fitted from the ESP of the charge densities of the ground state for the oxidized ($g_+$) and the neutral (g) P700 pair pigment $P_B$. Note that we assume localization of the positive charge at this pigment (*44–47*). $\Delta E_{\mathrm{P_A}}^{(\mathrm{CT})}$ contains short-range contributions to the electrochromic shift of $P_A$ arising from its proximity to $P_B$. The value of $\Delta E_{\mathrm{P_A}}^{(\mathrm{CT})}$ will be obtained from a fit of the experimental difference spectrum. The difference charges are rescaled by a constant factor to reproduce the experimental value of the difference dipole moment $\Delta\mu = 1$ D for Chl *a* (*56*). The same correction factor is used to rescale the difference charges of Chl *f*, for which our quantum chemical calculations obtain a difference dipole moment that is about half in magnitude compared to that obtained for Chl *a*. A temperature-dependent effective dielectric constant $\epsilon_{\mathrm{eff}}$, determined in earlier work (*57*) is applied. The dielectric constant was found to vary between $\epsilon_{\mathrm{eff}} = 2$ at cryogenic temperatures and $\epsilon_{\mathrm{eff}} = 8$ at room temperature .

In the calculation of optical spectra $\alpha(\omega)_{\mathrm{P}}$ and $\alpha(\omega)_{\mathrm{P+}}$ of PSI, we divide the photosystem into domains of strongly coupled pigments with strong intra-domain and weak inter-domain excitonic couplings. A pigment belongs to a certain exciton domain if it is coupled stronger than a certain cut-off value $V_c$ to at least one pigment of that domain. Exciton delocalization is only allowed within the exciton domains. In this way dynamic localization effects of the exciton wavefunction are taken into account implicitly (*57*).

The exact value of $V_c$ is difficult to evaluate, it should, however, be in the same order of magnitude as the local reorganization energy of the exciton-vibrational coupling. For the present calculations we used $V_c = 30\ \mathrm{cm}^{-1}$. The optical spectra $O(\omega)$ of PSI are then obtained as a sum over the optical spectra of the domains $O(\omega) = \sum_d O_d(\omega)$. The exciton Hamiltonian of every exciton domain is diagonalized separately revealing the eigen energies $\hbar\omega_{M_d}$ and the coefficients $c_{m_d}^{(M_d)}$ of the exciton states

$$|M_d\rangle = \sum_m c_{m_d}^{(M_d)} |m_d\rangle. \qquad (6)$$

The coefficients are used to calculate the transition dipole moments of the exciton states.

$$\boldsymbol{\mu}_{M_d} = \sum_{m_d} c_{m_d}^{(M_d)} \boldsymbol{\mu}_{m_d}, \qquad (7)$$

which determine the intensities of optical lines in the linear absorption spectrum.

$$\alpha_d(\omega) \sim \omega \sum_{M_d} \left|\boldsymbol{\mu}_{M_d}\right|^2 D_{M_d}(\omega) \qquad (8)$$

where $D_{M_d}(\omega)$ is the lineshape function obtained with a second-order cumulant expansion using partial ordering prescription (POP) and a Markov and secular approximation for the off-diagonal elements of the exciton-vibrational coupling in the exciton basis.

$$D_{M_d}(\omega) = \frac{1}{2\pi} \int_{-\infty}^{\infty} dt\, e^{i\left(\omega - \tilde{\omega}_{M_d}\right)t} e^{G_{M_d}(t) - G_{M_d}(0)} e^{-\frac{t}{\tau_{M_d}}} \qquad (9)$$

where the 0-0 transition frequency $\tilde{\omega}_{M_d}$ contains a renormalization by the exciton-vibrational coupling.

$$\tilde{\omega}_{M_d} = \omega_{M_d} - \frac{E_\lambda^{(M_d)}}{\hbar} + \sum_{K_d}^{K_d \neq M_d} \tilde{C}_{M_d K_d}^{(\mathrm{Im})}\left(\omega_{M_d K_d}\right) \qquad (10)$$

with $\omega_{M_d K_d} = \omega_{M_d} - \omega_{K_d}$, the reorganization energy of the $M_d$th exciton state.

$$E_\lambda^{(M_d)} = \sum_{m_d} \left(c_{m_d}^{(M_d)}\right)^4 E_\lambda^{(m_d)} \qquad (11)$$

which is related to the local reorganization energy $E_\lambda^{(m_d)}$ of the exciton of piment $m_d$ that is determined by the spectral density of the local exciton-vibrational coupling $J_{m_d}(\omega)$ that describes the modulation of the transition energy of site $m_d$.

$$E_\lambda^{(m_d)} = \int_0^\infty d\omega\, \hbar\omega\, J_{m_d}(\omega) \qquad (12)$$

$\tilde{C}_{M_d K_d}^{(\mathrm{Im})}(\omega)$ in Eq. 10 is related to the imaginary part of the half-sided Fourier transform of the correlation function of the electronic energy gap $C_{m_d}(t)$of the pigments.

$$\tilde{C}_{M_d K_d}(\omega) = \int_0^\infty dt e^{i\omega t} \sum_{m_d} \left(c_{m_d}^{(M_d)}\right)^2 \left(c_{m_d}^{(K_d)}\right)^2 C_{m_d}(t) \qquad (13)$$

obtained from the local spectral density $J_{m_d}(\omega)$ as:

$$C_{m_d}(t) = \int_0^\infty d\omega\, J_{m_d}(\omega)\omega^2 \left(\left(1 + n(\omega)\right)e^{-i\omega t} + n(\omega)e^{i\omega t}\right). \qquad (14)$$

The function $G_{M_d}(t)$ in the lineshape function in Eq. 9 describes the excitation of vibrational sidebands and is given as:

$$G_{M_d}(t) = \sum_{m_d} \left(c_{m_d}^{(M_d)}\right)^4 \int_0^\infty d\omega\, J_{m_d}(\omega) \left(\left(1 + n(\omega)\right)e^{-i\omega t} + n(\omega)e^{i\omega t}\right). \quad (15)$$

The inverse lifetime $\tau_M^{-1}$ of the $M$th exciton state in Eq. 9 is obtained from the real part of $\tilde{C}_{M_d K_d}(\omega)$.

$$\tau_{M_d}^{-1} = \sum_{K_d}^{K_d \neq M_d} \tilde{C}_{M_d K_d}^{(\mathrm{Re})}\left(\omega_{M_d K_d}\right) \quad (16)$$

and results in life-time broadening induced by exciton relaxation. We use the functional form of the spectral density extracted from fluorescence line narrowing spectra of B777 complexes (*58*). The integral coupling strength, that is the Huang-Rhys factor $S_{m_d} = \int_0^\infty d\omega\, J_{m_d}(\omega)$, has been inferred from the temperature dependence of the linear absorption spectrum of PSI, assuming a site-independent spectral density. The latter analysis resulted in $S_{m_d} = S = 0.6$ (*59*), which gives a reorganization energy $E_\lambda = 61$ $cm^{-1}$. In order to take into account, the coupling to CT states and to improve the description of the $P700^+$ *minus* P700 difference spectrum, the Huang-Rhys factors of the RC pigments are allowed to differ from that of the antenna pigments.

Besides homogeneous broadening (described above), inhomogeneous broadening arising from slow conformational motion of the protein, needs to be taken into account. We assume a Gaussian distribution functions for the local transition energies of the pigments that are centered around the mean values (site energies) and have a certain width that may also depend on the site. Many realizations of site energies are generated randomly from these distribution functions and the homogeneous spectra calculated for these realizations are averaged to obtain the inhomogeneous absorption spectrum. For the calculation of the $P^+$ absorption spectrum of PSI, we replace the site energies $E_m(\mathrm{P})$ used for the calculation of the PSI absorption spectrum by $E_m(\mathrm{P}^+)$ that contain the electrochromic shift induced by $\mathrm{P}^+$ (Eqs. 4 and 5), and we switch off all excitonic couplings involving $P_B$, where the positive charge is assumed to be localized. In addition, we consider the featureless broad absorption spectrum of the oxidized $P_B$, which is estimated from experimental data (*60*) on the absorption of Chl *a* and its cation Chl $a^+$ as described in the following.

Absorption spectrum of Chl $a^+$

For the Chl $a^+$ our quantum chemical calculations find 5 excited states approximately in the usual absorption region of Chl *a*, but somewhat red-shifted. Summing up their dipole strengths gives about 3.4 D in comparison to 5.13 D obtained for the $Q_y$ transition of Chl *a*. Chauvet et al (*60*) measured the absorption spectra of both, Chl *a* and Chl $a^+$, showing the latter has a broad, red-shifted absorption band, which reaches from 700 nm to 900 nm. Integration over this broad band reveals an effective transition dipole moment that reaches 80 % of that obtained for the $Q_y$ transition by integrating over the low-energy absorption band of Chl *a*. This ratio is in qualitative agreement with the quantum chemical calculation reported above. In the calculation of the $P^+$-P difference spectrum, we model the absorption of $P_B^+$ by a band with a transition dipole moment that is 0.8 times that of Chl *a*, only red-shifted to 720 nm, and with a much smaller width (50 nm instead of 200 nm) than measured on Chl $a^+$ in solution. This width is obtained by increasing the inhomogeneous width (Michael: give value) and the homogeneous width (Huang-Rhys factor 5.2). At present these values are just obtained from a fit of the broad positive experimental band occurring between 720 and 740 nm in the difference spectrum. Note, however, that this band is rather independent on the remaining much sharper features of the difference spectrum used to check the assignment of Chl *f*, as will be discussed in more detail below.

Quantum chemical calculations of charge- and transition densities of Chl *a* and Chl *f*

Isolated Chl *a* and Chl *f* were geometry-optimized with density functional theory using the CAM-B3LYP exchange correlation (XC) functional and the polarization consistent basis set pcseg-1. Charge densities of the ground state and the excited state of Chl *a*, Chl $a^+$ and Chl *f,* as the transition densities and transition dipole moments, were calculated using (time-dependent) density functional theory (TDDFT). For those quantum chemical **calculations,** the wB-97X-V XC-functional and the pcseg-1 basis set were used. The electrostatic potential of the charge and transition densities were fitted by atomic partial charges centred on the heavy atoms of the chlorophylls, CHELP-BOW (*61*), following the TrEsp  and CDC methods (Adolphs et al., 2008; M. E. Madjet et al., 2006). These atomic partial charges were rescaled to match experimental values for transition and difference dipole moments, as described above.

**Supplementary text S1: Chl *f* site assignments in common with literature**

Chl A21

The ESP map of A21 indicates that it is a Chl *f*, with its formyl group H-bonded (2.83 Å) by the backbone amide of PsaA2 $Leu^{333}$(fig. S2A). Statistical analysis of the local ESP around the C2 substituent confirms the presence of a formyl group (fig. S2B, S2C). $Leu^{333}$ is part of a loop insertion of 8 amino acids specific to far-red PsaA2 (fig. S2D). In addition, the insertion loop contains a conserved $Met^{332}$, located with the sulfur atom 3.45 Å from the formyl oxygen, suggesting a possible stabilising interaction. A water molecule, part of a group of water molecules forming a H-bond network, also involving PsaA2 $Gln^{323}$, acts as the axial ligand for the $Mg^{2+}$. In WL-PSI, A21 is coordinated by PsaA $His^{332}$ (*41*). The insertion disrupts this site by changing the A21 ligand from His to a water and by creating the stabilising environment for the formyl group. A21 was previously identified as a Chl *f* candidate based on i) the loss of the His and the presence of water as the axial ligand, and ii) the presence of a potential H-bond donor. The ESP map from *Synechococcus sp.* PCC 7335 and *H. hongdechloris* also showed indications of the expected formyl group (Gisriel et al., 2021; Kato et al., 2020).

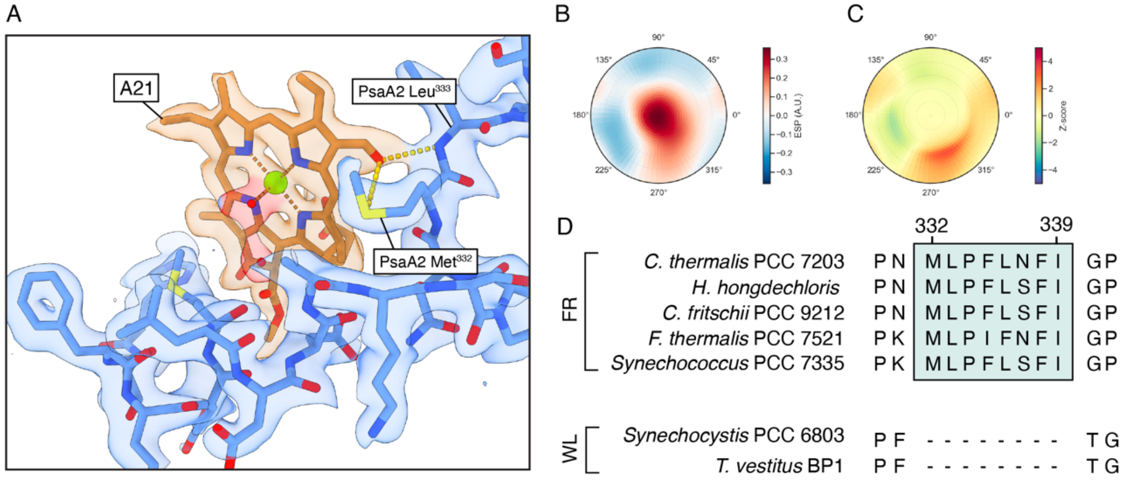


**fig. S2 Chlorophyll A21**

A) Chemical environment of A21: the Chl density is in orange, the density of the amino acid residues from PsaA2 in blue, the density of water molecules in red. H-bonds are indicated as dotted lines. The C2 formyl group is within H-bonding (2.83Å) distance to the peptide bond amide group of PsaA2 Leu$^{333}$ and forming a favorable interaction (3.45Å) with PsaA2 Met$^{332}$. B) ESP polar plot indicating the presence of an increased potential around the C2 substituent at an angle of ~270°. C) Z-score polar plot indicating the presence of an increased potential around the C2 substituent at an angle of ~270°. D) Multiple sequence alignment of FR and WL PsaA, showing the far-red specific loop insertion of 8 amino acids.

Chl A23

The A23 Chl *f* is identified here along with its H-bonding ligand, PsaA2 $Gln^{365}$ (fig. S3A, S3B, S3C). The same assignment was made in *H. hongdechloris* based on structural and phylogenetic arguments (*9*, *10*). It was noted that among Chl *f* producing species, a small majority lack the Gln and instead have the non-H-bonding Met, leading to the assumption that in the presence of PsaA2 $Met^{365}$, the site binds a Chl *a* (*9*, *10*).

Other conserved changes are identified in the helix that coordinates A23 in the species that have Gln H-bonded to the Chl *f*. These changes may be important for the insertion of a Chl *f* into this site (fig. S3D). PsaA2 $Thr^{368}$ H-bonds the oxygen of PsaA2 $Gln^{365}$ keeping the amide group in a position that favours the formation of the H-bond with the C2 formyl oxygen. At the same time, two additional changes, PsaA2 $Asn^{358}$ and PsaA2 $His^{362}$, move the water that coordinates the $Mg^{2+}$ of A23, shifting the chlorophyll to a position that optimises the interaction of the C2 formyl group with PsaA2 $Gln^{365}$. In addition, a series of partially conserved changes can be identified in the species that have the PsaA2 $Gln^{365}$. PsaA2 $Met^{196}$ is a Leucine in other species, and PsaA2 $Trp^{192}$ is a Valine in most of the other species, these amino acids could interact with the π-electrons of the chlorophyll ring, affecting the absorption of A23.

Methionine can form stable interactions with carbonyl oxygens in proteins, where the sulfur is thought to act as an electrophile to electron-rich atoms acting as nucleophiles, such as the formyl oxygen of Chl *f* (*62*). The effect of this interaction, on the spectroscopic properties of a chlorophyll, is expected to be different to that of an H-bond. It is also likely that this interaction is weaker at stabilising the binding of Chl *f*, compared with an H-bond, in agreement with the presence of Chl *a* instead of Chl *f*, as mentioned above. Phylogenetic comparison of FR and WL-PSI shows that in FR-PSI, two options with near equal frequency are found at this location, Met or Gln, while in WL-PSI two different options still with near equal frequency are found, Met or Leu. Leucine, a residue incapable of interacting and stabilise a Chl *f* in position A23, is never found in FR-PSI, this might suggest that is some species in which PsaA2 $Met^{365}$ is present, A23 is a Chl *f* or that the ancestor of these FR-species had a methionine in this position.

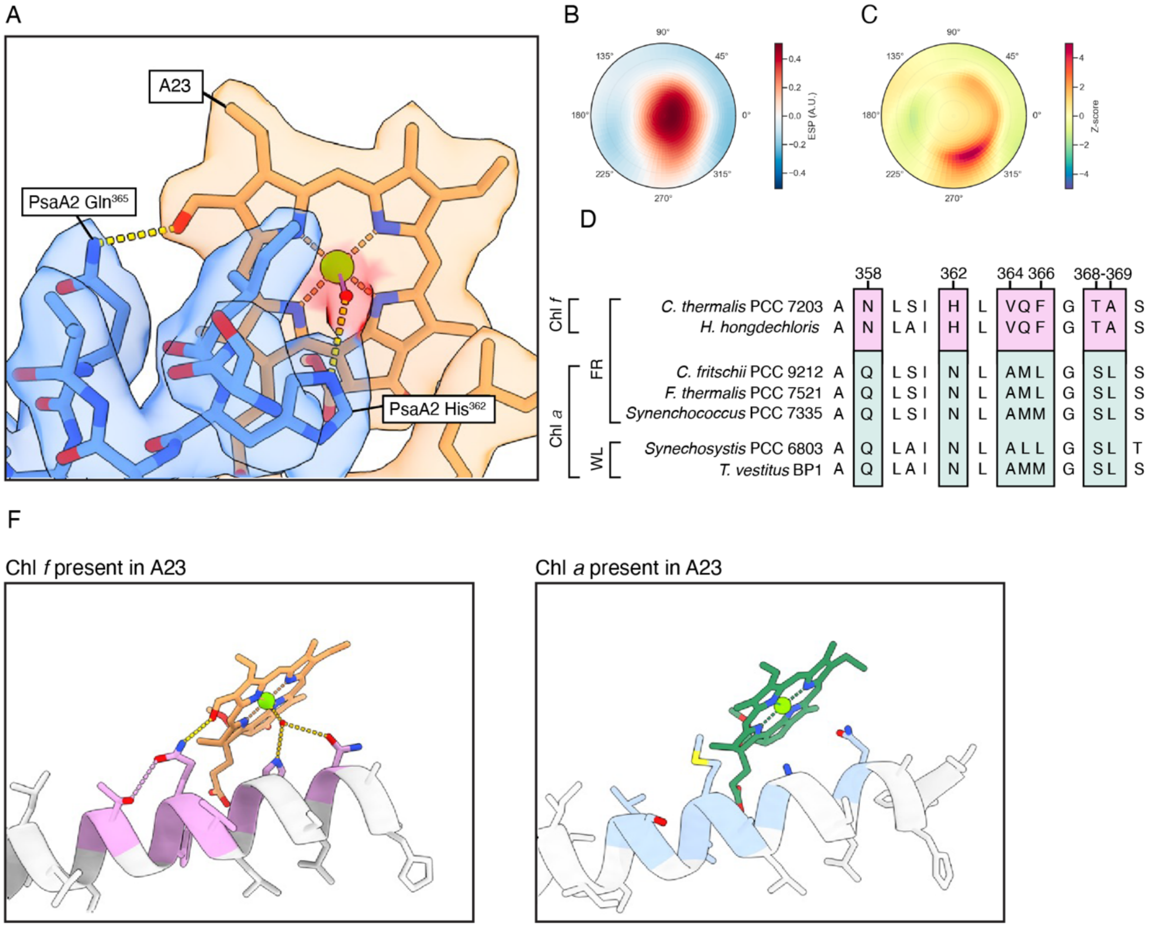


**fig. S3 Chlorophyll A23**

(A) Chemical environment of A23. The Chl density is in orange, the density of the amino acid residues in blue, the density of water molecules in red. H-bonds are indicated as dotted lines. (B) ESP polar plot indicating the presence of an increased potential around the C2 substituent. (C) Z-score polar plot indicating the presence of an increased potential around the C2 substituent. (D) Multiple sequence alignment of FR and WL PsaA, showing the differences that allow the binding of a Chl *f* at the A23 site. In pink, changes associated with the presence of a Chl *f*, and in light blue the equivalent amino acids in WL-PSI and in those FR species that appear to lack Chl *f* in this site. (F) Model of the interactions between PsaA2 and Chl A23 in *C. thermalis* with Chl *f* and *F. thermalis* with Chl *a.*

Chl B07

The ESP map indicates an H-bond from the back-bone amide of PsaB2 $Ala^{94}$ to the C2 formyl group of Chl *f* in the B07 site (fig. S4A, S4B, S4C). PsaB2 $Ala^{94}$ is present in most far-red species instead of a conserved proline in WL-PSI. The $Mg^{2+}$ of the Chl *f* is coordinated by the amide of PsaB2 $Gln^{95}$ (fig. S4D). B07 was identified as a likely candidate for Chl *f* in earlier work, based on sequence-based modelling indicating a non-His axial ligand (*4*) and cryo-EM data (*8–11*). In *Synechococcus* sp. PCC 7335 (*11*), the presence of the WL specific Pro suggests that the formyl group is oriented differently, a water molecule is instead present in the cryo-EM structure of FR-PSI of *Synechococcus* sp. PCC 7335 together with an increase ESP potential from the C2 formyl substituent.

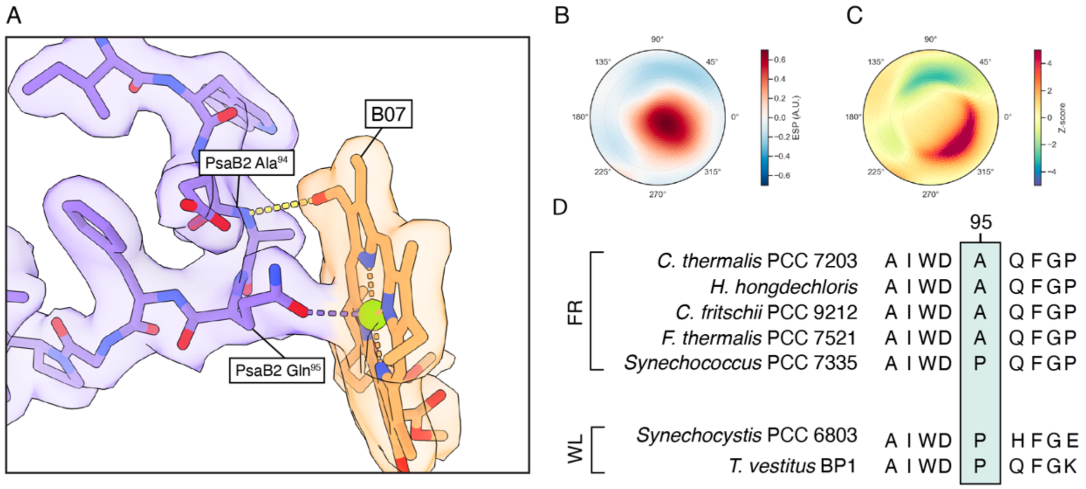


**fig. S4 Chlorophyll B07**

(A) ESP map and atomic model around the B07 site. The Chl density is in orange, the density of the amino acid residues in PsaB2 is in violet. H-bonds are indicated as dotted lines. (B) ESP polar plot indicating the presence of an increased potential around the C2 substituent. (C) Z-score polar plot indicating the presence of an increased potential around the C2 substituent. (D) Multiple sequence alignment of FR and WL PsaB, showing Pro to Ala substitution in position 95 that is specific to the far-red sequence, although not present in all of them.

Chl B30

The ESP map shows an H-bond to the formyl group of B30 from $Tyr^{38}$, a conserved change present in PsaJ2 (fig. S5A, S5D). Local analysis of the ESP results in a clear single peak oriented towards the PsaJ2 $Tyr^{38}$ sidechain (fig. S5B, S5C). $Tyr^{38}$ is part of the C-terminal extension of PsaJ2 that displaces three Chl *a* molecules and a carotenoid from this region in the canonical WL structure (CLA 1301,1302,1303 and BCR 1306 in (*41*)). B30 also shows an unusual ε-His ligand from PsaB2 $His^{443}$. Calculations suggest that ε-His coordinated chlorophylls have slightly lower site-energies than those with the more common δ-His ligand (*63*). Chl B30 was previously identified as a Chl *f* candidate given the potential H-bond from the conserved FR-specific PsaJ2 $Tyr^{38}$ (*28*). When the cone scan was applied, indications for a distribution between two possible orientations were reported in *F. thermalis* , but this evidence was lacking in *H. hongdechloris* as the PsaJ2 subunit was absent from the structure (*9*). The lower resolution in *F. thermalis*, the two possible positions for the proposed H-bond, and the anomalously long His-$Mg^{2+}$ distance, all indicate that in the *F. thermalis* FR-PSI this site was perturbed. Neither the ESP map nor the cone scan for *Synechococcus* sp. PCC 7335 showed evidence for the formyl in B30 (*11*). This was interpreted as low occupancy of the site, perhaps due to loss of PsaJ2 as seen in *H. hongdechloris* in both FR- and WL-PSI. Here, in intact FR-PSI map with higher resolution, this site is fully occupied.

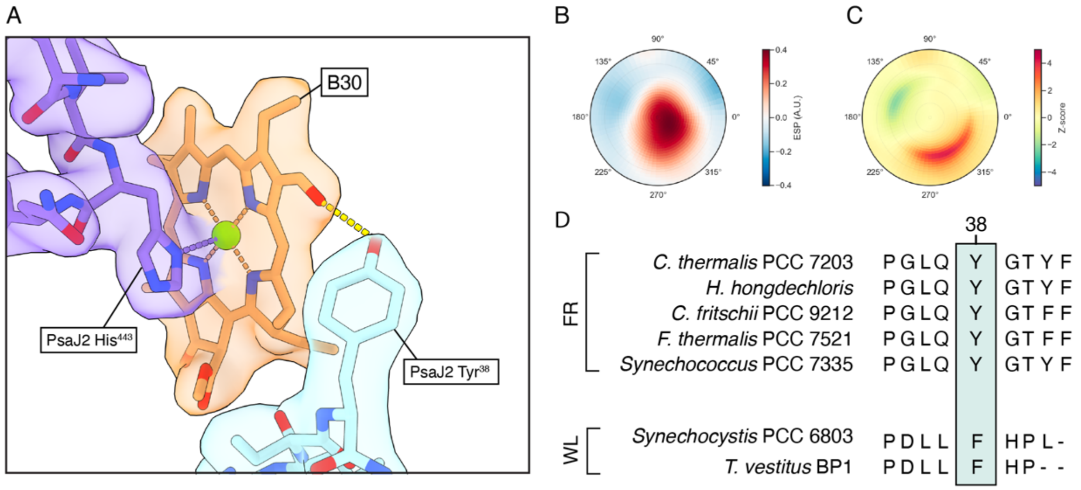


**fig. S5 Chlorophyll B30**

(A) ESP map and atomic model around the B30 site. The Chl density is in orange, the density of the amino acid residues from PsbB2 in violet and those from PsaJ2 in cyan. H-bonds are indicated as dotted lines. (B) ESP polar plot indicating the presence of an increased potential around the C2 substituent. (C) Z-score polar plot indicating the presence of an increased potential around the C2 substituent. (D) Multiple sequence alignment of FR and WL PsaJ, showing the far-red specific PsaA2 Tyr$^{38}$.

Chl B37

An H-bond from the backbone amide of PsaB $Gly^{697}$ to the formyl oxygen of B37 (fig. S6A) is evident in the ESP polar plot of the C2 substituent (fig. S6B, S6C). PsaB2 $Gly^{697}$ is specific for FR-PSI, replacing either Val or Ile in PsaB of WL-PSI (fig. S6D). The $Mg^{2+}$ of B37 has water as its axial ligand. Based on these characteristics, B37 is confirmed as being a Chl *f*, in agreement with previous assignments (*4*, *8*, *9*, *13*).

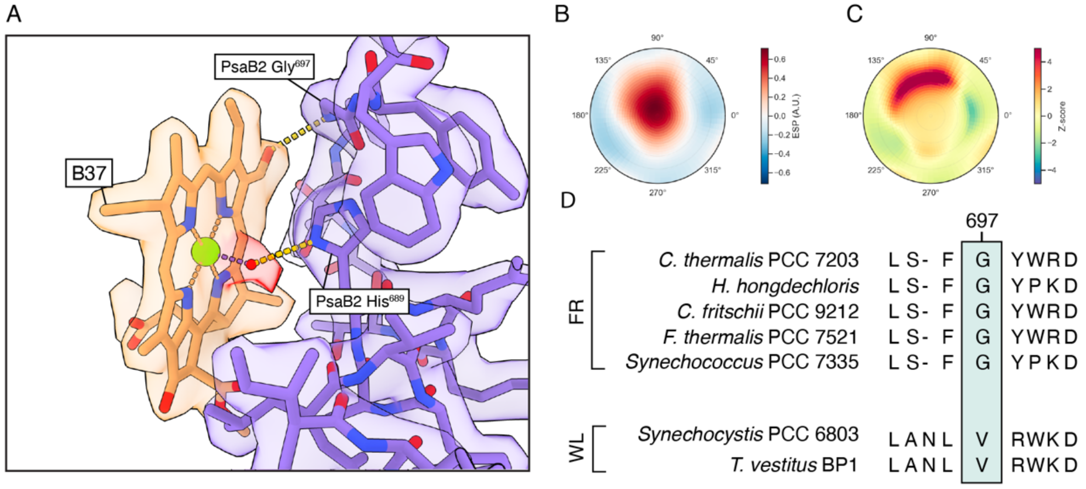


**fig. S6 Chlorophyll B37**

(A) ESP map and atomic model around the B37 site. The Chl density is in orange, the density of the amino acid residues in violet and the density from water in red. H-bonds are indicated as dotted lines. (B) ESP polar plot indicating the presence of an increased potential around the C2 substituent. (C) Z-score polar plot indicating the presence of an increased potential around the C2 substituent. (D) Multiple sequence alignment of FR and WL PsaB, showing the FR-specific Gly in position 697.

Chl B38

Together with B37, B38 forms the low energy Chl *f* coupled pair absorbing at 800 nm (*4*, *8*, *10*, *13*). The central $Mg^{2+}$ of B38 is coordinated by a water that forms a distal H-bond with PsaB2 $Asp^{701}$, but its formyl oxygen lacks a hydrogen bond partner (fig. S7A, S7D). Nevertheless, a positive feature, with peak Z-score > 3 is present in the ESP polar plot of the C2 substituent. The torsion angles of the feature suggests that the formyl group is parallel to PsaB2 $Trp^{22}$, a residue that is also conserved in PsaB in WL-PSI. Compared to other sites, B38 presents a lower intensity of the signal, reflecting the weak character of this interaction, and once again confirming that higher electronegativity and a broader spread of torsion angles can diminish the formyl oxygen's signal. Tros et al. and Gisriel et al. (*11*, *13*) pointed to a conserved change in some FR-PSI sequences, where PsaB $Trp^{22}$ is replaced with a Phe, which being smaller, allows the insertion of a water molecule to bridge the formyl oxygen of Chl B38 to the carbonyl backbone of $Thr^{18}$. Nevertheless, this change is not conserved in all FR-PSI sequences. Indeed, it remains a Trp in *C. thermalis* PCC 7203 and there is no density indicating that a water molecule is present in our structure.

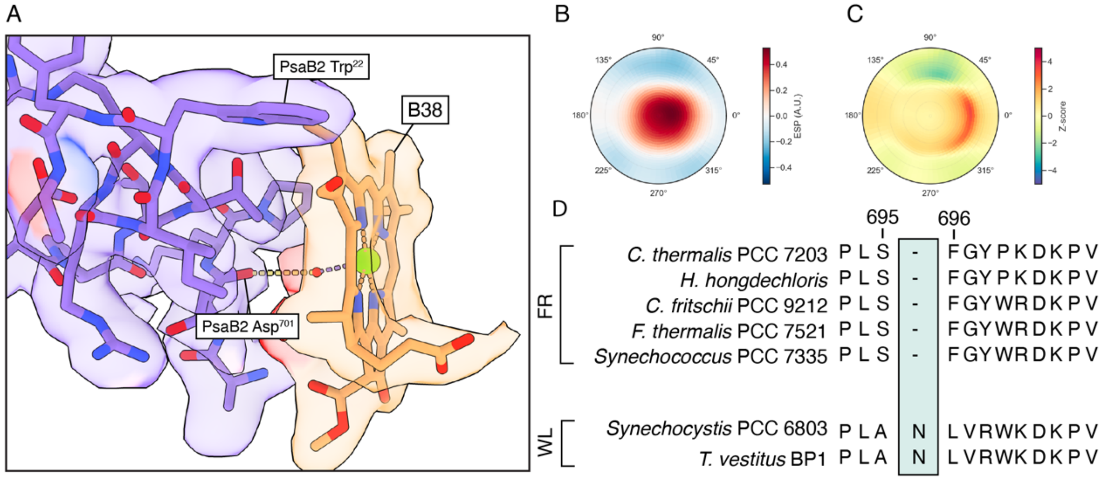


**Fig. S7 Chlorophyll B38**

(A) ESP map and atomic model around the B38 site. The Chl density is in orange, the density of the amino acid residues in violet and the density from water in red. H-bonds are indicated as dotted lines. (B) ESP polar plot indicating the presence of an increased potential around the C2 substituent. (C) Z-score polar plot indicating the presence of an increased potential around the C2 substituent. (D) Multiple sequence alignment of FR and WL PsaB.

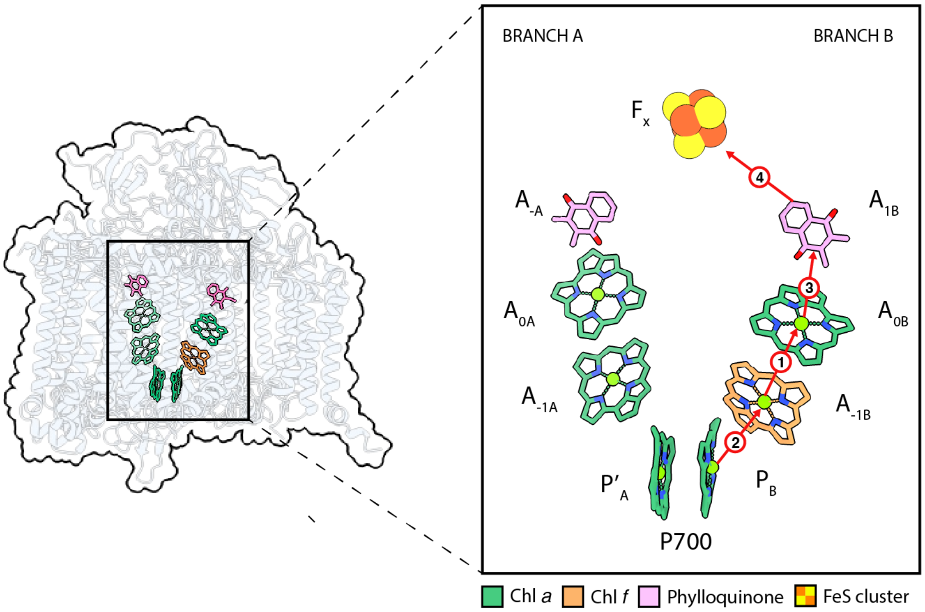


**fig. S8 FR-PSI electron transfer cofactors.**
The cofactors of the two branches of electron transfer are shown together with the $F_X$ cluster. Chl *f* is shown in orange in the $A_{-1B}$ position. The red arrows represent the suggested steps of electron transfer upon charge separation.

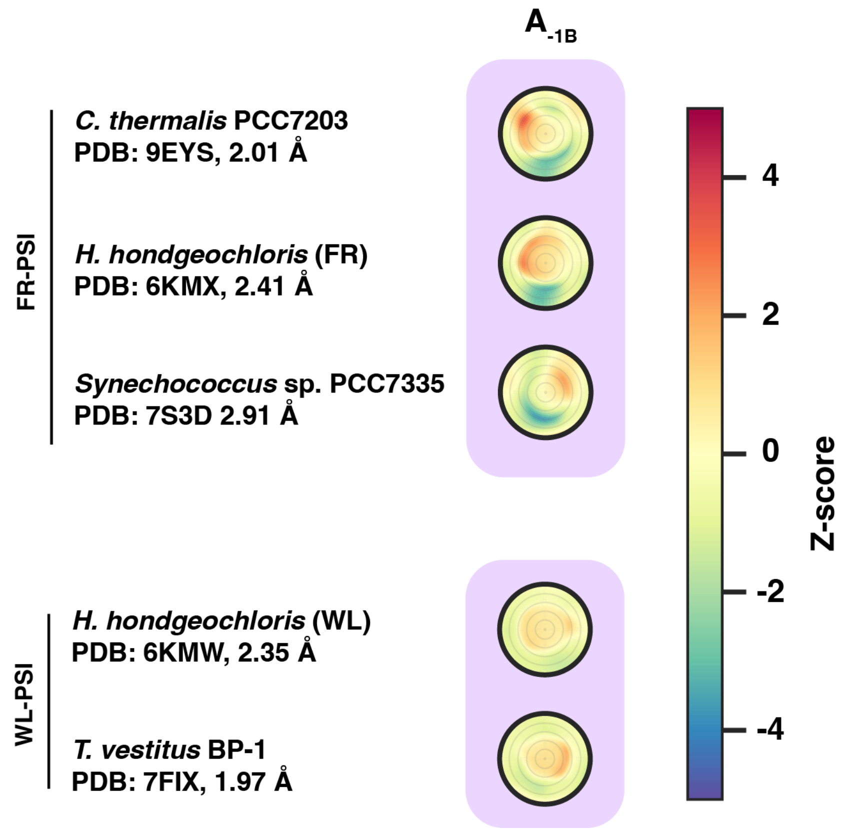


**fig. S9 Polar plots of the ESP at the C2 position for the $A_{-1B}$ chlorophylls of FR-PSI and WL-PSI from different species.**

Comparison of plots of ESP of the C2 substituent on the $A_{-1B}$ chlorophylls calculated using the high-resolution structures for FR-PSI and WL-PSI available from the PDB. The plots show the Z-scores colored according to the scale bar, with statistically significant positive and negative regions of ESP present only in the FR-PSI.

### Supplementary Text S2: DFT calculation of the $A_{-1B}$ chemical environment

To identify and distinguish the origins of the electrostatic effects contributing to the increased negative ESP features observed around the C2 formyl of Chl *f* in the $A_{-1B}$ site DFT calculations were performed. These calculations were done on the $A_{-1B}$ and $A_{0B}$ chlorophylls and the surrounding protein, as described in the Materials and Methods, using the structural model with and without an additional optimization step. In the following, the qualitative results apply to both optimized and non-optimized structures.

With no H-bond to the formyl group, the negatively polarized formyl oxygen remains un-neutralized, exhibiting a negative well of ESP (fig. S10B, S10D). This feature is present irrespective of the presence of PsaA2 $Phe^{459}/Cys^{463}$. When PsaA2 $Phe^{459}/Cys^{463}$ is present, the negative ESP well is spatially confined to the region between the Chl *f* and the PsaA2 $Phe^{459}$ (fig. S10B), and the depth of the well is estimated to decrease from -0.0667 to -0.0432 atomic units (~0.640 eV). With Chl *a* in the far-red $A_{-1B}$ site, its C2 methyl group is adjacent to PsaA $Phe^{459}/Cys^{463}$, but there is no negative well between the Chl *a* and the PsaA2 $Phe^{459}$ (S10A). Thus, the presence of the negative ESP feature in the cryo-EM is additional evidence of Chl *f* being in the $A_{-1B}$ position.

The presence of a formyl group at the C2 site of Chl *f* leads to a redistribution of ESP along the y-axis of the Chl $A_{-1B}$ (fig. S10E), and this is directed by the interaction of the C2 formyl bond and the π - electrons of PsaA2 $Phe^{459}$. This interaction together with the fact that alignment of the formyl C=O bond with the chlorophyll plane is the lowest energy configuration, contribute to the preferred location of the formyl oxygen and therefore to the location of its associated negative ESP, as observed experimentally. The negative feature at ~270° observed in the Z-score polar plot (fig. 2C) arises from the intersection of this negative well with the conical surface of the scan. The negative feature is augmented by the subtraction of the average C7 methyl environment, which includes a ring of positive ESP at the averaged methyl hydrogen radius (see Quantification and statistical analysis of the electrostatic potential).

The presence of the negative ESP feature in the region of the C2 formyl oxygen suggests that the sulfhydryl-π-formyl motif has functional and evolutionary significance in tuning the binding and the redox potential of the Chl *f* in the $A_{-1B}$ site. Chl $f^{-\bullet}$ and Chl $f^{*}$ are less reducing than Chl $a^{-\bullet}$ Chl $a^{*}$ and so their redox potentials must be lowered if Chl *f* is to play redox roles in PSI like those of Chl *a* (*25*). These chemical requirements explain why there is no conserved H-bonding group to facilitate its binding, nor to neutralize the negatively polarized formyl oxygen and thereby decreasing its reducing power. This also explains why the positive ESP signal arising from the C2 formyl oxygen of the Chl *f* in the $A_{-1B}$ site required much better resolution for its detection.

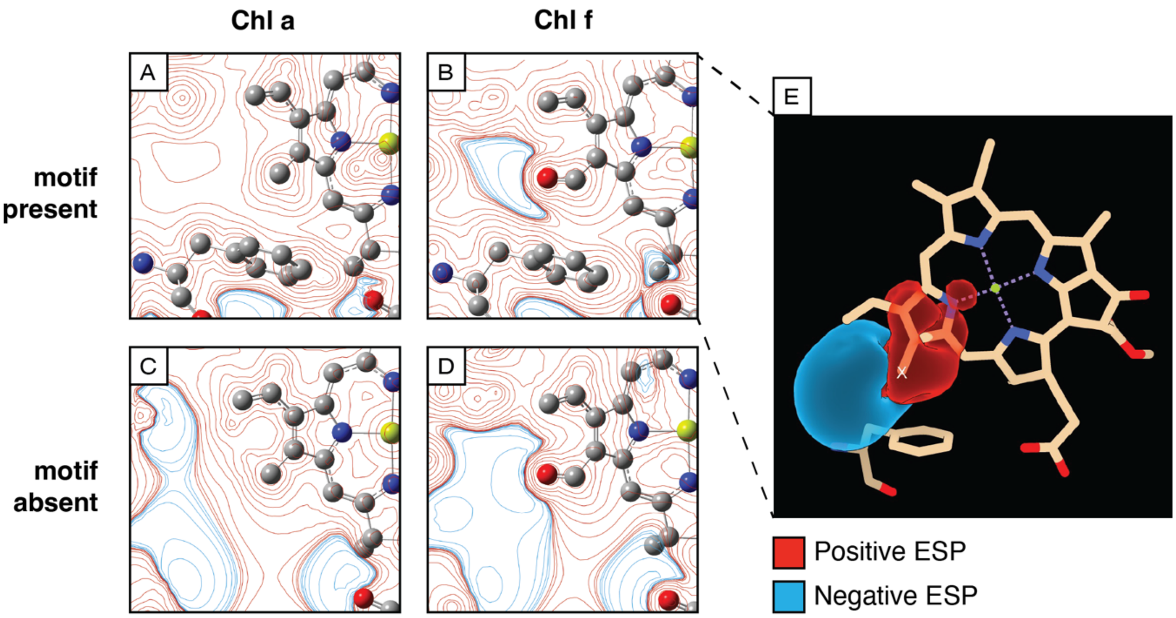


**fig. S10: DFT calculations of the $A_{-1B}$ environment**

DFT-calculated ESP contours in a plane defined by the formyl group (C=O) at the C2 position of Chl $A_{-1B}$ and the γ-carbon of PsaA Phe[459]. Contours are shown for ESP values of ±0.001, ±0.002. ±0.004, ±0.008, ±0.02, ±0.04, +0.08, +0.02, +0.4, +0.8, +2 (all atomic units). Red indicates positive ESP, whilst blue indicates negative ESP. A) Chl *a* in the $A_{-1B}$ site with PsaA Phe[459]/Cys[463] present. B) Chl *f* in the $A_{-1B}$ site with PsaA Phe[459]/Cys[463] present. C) Chl *a* in the $A_{-1B}$ site with PsaA Phe[459]/Cys[463] absent. D) A Chl *f* in the $A_{-1B}$ site with PsaA Phe[459]/Cys[463] absent. The negative ESP on the left-side of fig. S10C arises from an oxygen of the Chl *a* in the $A_{0B}$ site, which is above the plane of the map. Panel A-D also show $ChlA_{-1B}$ and PsaA Phe[459]/Cys[463] as ball and stick models. The Chl $A_{0B}$ and the other amino acids adjacent to the C2 carbon were included in the calculation but are not shown in the figure.

E) ΔESP map obtained by subtracting the Chl *a* ESP map (S20A, methyl at X) from the Chl *f* ESP map (S20B, C2' of formyl at X). The ΔESP map is shown as iso-surfaces at ±0.01 atomic units, where all the ΔESP values more negative than -0.01 a.u. are contained within the blue (negative change) surface and all values greater than 0.01 a.u. are contained within the red (positive change) surface. The figure also shows Chl $A_{-1B}$ and PsaA Phe[459] as a stick model. The Chl $A_{0B}$ and the other amino acids adjacent to the C2 carbon were included in the calculation but are not shown in the figure. The structural model used for these calculations were based on the cryo-EM map directly without optimization.

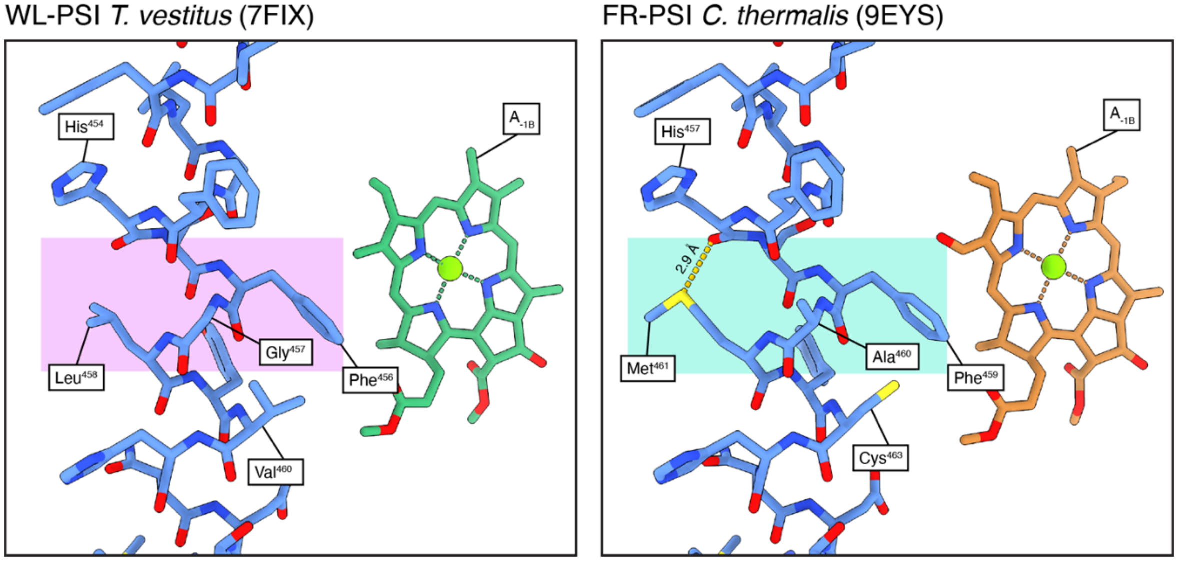


**fig. S11 Structural comparison of Helix N PsaA and PsaA2 in the region $A_{-1B}$**

The panels show a structural comparison of the PsaA and PsaA2 in high resolution structures of WL-PSI (7FIX) and FR-PSI (9EYS) respectively. In both forms of PSI, the helix turn that contains the $Phe^{456}$/$Phe^{459}$ facing Chl $A_{-1B}$ is distorted, due to the pull exerted by the central $Mg^{2+}$ of Chl A31 (not shown) on $His^{454}$/$His^{457}$. In the WL-PSI this results in the backbone carbonyl of PsaA $His^{454}$ being too distant from the amide backbone of PsaA $Leu^{458}$ to be involved in an α-helical H-bond (~5Å). While in WL-PSI this specific turn is likely to be flexible due to the lack of the α-helical H-bond and the presence of PsaA $Gly^{457}$, in the FR-PSI, two of the conserved changes, PsaA2 $Ala^{460}$ and PsaA2 $Met^{461}$, seem to stabilize the helix turn. PsaA2 $Met^{461}$ interacts with the carbonyl of PsaA2 $His^{457}$, re-establishing a connection between the two helix turns. This could have consequences on the rigidity of this portion of the protein, especially during the electron transfer process, and for holding in place the two residues, PsaA2 $Phe^{459}$ and PsaA2 $Cys^{463}$, which are proposed to be important for the function of Chl *f* $A_{-1B}$ as the primary donor. For WL-PSI the destabilized helix turn is highlighted in pink and for FR-PSI the same region with the additional stabilizing interaction, is highlighted in cyan.

**Supplementary Text S3: Electrochromic shifts data and discussion**

In general, the electrochromic shifts $\Delta E_m$ (Eq. 5) of the antenna pigments are small because of the large distance to the location of the charge, $P_B^+$, in the P700 pair. We identify three antenna chlorophylls with an electrochromic shift that is sufficiently large to influence the $P^+$-P difference spectrum directly (and not only indirectly via excitonic coupling modulating the delocalization). These are the two linker pigments A40 and B39 and the antenna Chl B24, which is closest to $P_B$ resulting in the largest electrochromic shift, for both pigment types Chl *a* and Chl *f* (table S1).

| *m* | Chl *a* $\Delta E_m$ (cm$^{-1}$) | Chl *f* $\Delta E_m$ (cm$^{-1}$) |
|---|---|---|
| A40 | -23.36 | -10.51 |
| B39 | -30.80 | -13.86 |
| B24 | -45.42 | -20.44 |

**table S1:** Electrochromic shifts of Chl *a* and Chl *f* in the 3 antenna binding sites most susceptible to being shifted by $P_B^+$.

| *m* | $\Delta E_m$ (cm$^{-1}$) | *m* | $\Delta E_m$ (cm$^{-1}$) |
|---|---|---|---|
| $P_A$ | -44.58 | $P_B$ | - |
| $A_{-1A}$ | -86.97 | $A_{-1B}$ | 9.03 |
| $A_{0A}$ | -4.42 | $A_{0B}$ | -27.08 |

**table S2**: Electrochromic shifts of Chl *a* in different binding sites in the reaction center.

| *m* | $\Delta E_m$ (cm$^{-1}$) | *m* | $\Delta E_m$ (cm$^{-1}$) |
|---|---|---|---|
| $P_A$ | 39.71 | $P_B$ | - |

| | | | |
|---|---|---|---|
| $A_{-1A}$ | -69.25 | $A_{-1B}$ | -45.88 |
| $A_{0A}$ | 7.05 | $A_{0B}$ | -5.09 |

**table S3:** Same as in Table 2 but for Chl *f*.

Tables S2 and S3 show the electrochromic shifts of the reaction center pigments. Interestingly, although the difference dipole moment for Chl *f* is only half in magnitude compared to that of Chl *a*, the electrochromic shifts in the reaction center are of similar magnitude or even larger than for Chl *a*. Especially the two closest neighbors of $P_B$, $P_A$ and $A_{-1B}$, show a shift three times larger indicating clear differences of the electrostatic potential for Chl *a* and Chl *f* at very short distances. We would also like to emphasize that the sign of the electrochromic shift at $A_{-1B}$ switches between Chl *a* and Chl *f*. Applying a dipole approximation to Chl *a* results in an increase of the electrochromic shift at $A_{-1B}$ from 9 $cm^{-1}$ (table S2) to 48 $cm^{-1}$ (table S6), whereas the dipole approximation for Chl *f* changes the electrochromic shift from -46 $cm^{-1}$ (table S2) to -8 $cm^{-1}$ (table S7).

These results show that the more sophisticated charge density coupling (CDC) method is necessary for an accurate description of the electrochromic shifts at short distances, particularly for Chl *f*. For the three antenna pigments in table S1, we find electrochromic shifts of similar magnitude as those obtained in the reaction center if Chl *a* is considered. However, when Chl *f* is present at those specific sites, the electrochromic shifts become much smaller, indicating that at these distances the dipole contribution dominates and the smaller difference dipole moment of Chl *f* is the decisive factor.

In table S4 the calculated electrochromic shifts of the antenna Chl *f* pigments as proposed by the present cryo-EM structural model are listed. All the electrochromic shifts are small with the largest value still being smaller than 10 $cm^{-1}$ in magnitude reflecting the larger distance of these pigments to the reaction center. Based on these calculations it can be ruled out that the main peaks at ~750 nm in the $P^+$- P difference spectrum are caused by these Chl *f* antenna pigments, as will be shown in detail below. Among the Chl *f* pigments, B30 and B07 exhibit the largest electrochromic shift magnitudes. As discussed in the main text and below, these pigments are candidates to explain the small but distinct bleaching around 762 nm in the $P^+$- P difference spectrum.

| $m$ | $\Delta E_m$ (cm$^{-1}$) |
|---|---|
| A21 | 1.60 |
| A23 | -0.86 |
| B07 | 9.61 |
| B30 | 7.18 |
| B37 | 0.88 |
| B38 | -1.13 |
| L03 | -2.77 |

**table S4:** Electrochromic shifts of the Chl *f* antenna pigments identified in the present cryo-EM model structure.

Electrochromic shifts obtained using the point-dipole approximation.

For comparison, we investigated the electrochromic shifts in the PSI reaction center calculated within the point-dipole approximation (PDA). In Table 5 the PDA as used in Langley et al. in (*48*) is used, where the difference dipole moment is assumed to be directed along the $N_B$-$N_D$ axis.

| $m$ | $\Delta E_m$ (cm$^{-1}$) | $m$ | $\Delta E_m$ (cm$^{-1}$) |
|---|---|---|---|
| $P_A$ | -249.53 | $P_B$ | - |
| $A_{-1A}$ | 70.25 | $A_{-1B}$ | 72.84 |
| $A_{0A}$ | -22.74 | $A_{0B}$ | -27.04 |

**Table S5**: Electrochromic shifts of RC pigments calculated within the point-dipole approximation. The difference dipole moment is assumed to be directed along the $N_B$-$N_D$ axis and a magnitude of 1 D (Chl *a*) is chosen. The corresponding shifts for Chl *f* are approximately half of the magnitude these values due to the smaller difference dipole of Chl *f*.

In these calculations we assumed a magnitude of 1D for the difference dipole moment, corresponding to Chl *a* (*56*). For Chl *f* , according to our quantum chemical calculations, this magnitude is about half, such that the resulting electrochromic shifts for Chl *f* are half the values given in Table S5. The electrochromic shift of $P_A$ is significantly larger than that obtained with the CDC method (tables S2 and S3) and inverted in sign. As already pointed out above, the electrochromic shifts in the positions $A_{-1A}$ and $A_{-1B}$ are also sign-inverted with respect to the CDC values of Chl *f* in table S7. In PDA both electrochromic shifts are positive. This result led to the conclusion in Langley et al., 2022 that Chl *f* cannot be situated at these binding sites. The present model, however, shows that one needs to go beyond the PDA for obtaining the correct sign of the electrochromic shifts of Chl *f* at $A_{-1A}$ and $A_{-1B}$.

Improved point-dipole approximation

The PDA can be improved by taking into account the direction of the difference dipole moment resulting from the quantum chemical calculations instead of assuming an orientation of the difference dipole along the $N_B$-$N_D$ axis. The results for this improved PDA are shown in tables S6 and S7 for Chl *a* and Chl *f*, respectively. Whereas for Chl *a,* the two dipole approximations give similar results, in case of Chl *f* qualitative differences are obtained. The differences between tables S5 and S7 are caused by the smaller magnitude (which approximately reduces the electrochromic shift by 50%) and the 76° rotation of the difference dipole moment of Chl *f* with respect to the $N_B$-$N_D$ axis, whereas for Chl *a* this angle is only 5° (fig. S11). In particular, the electrochromic shift of Chl *f* at the $A_{-1A}$ and $A_{-1B}$ sites is still an order of magnitude smaller than that obtained with the more accurate CDC method (table S3).

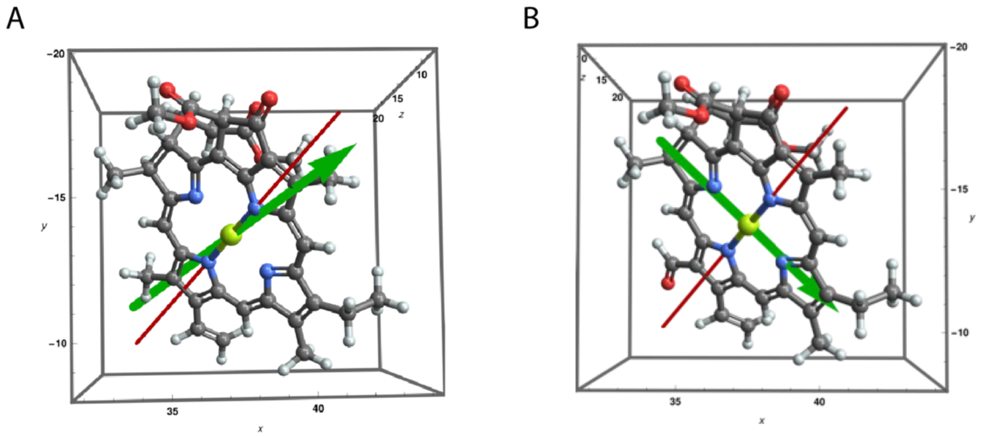


**Fig. S12: Difference dipole moment direction for Chl *a* and Chl *f***

Difference dipole moment direction for (A) Chl *a* (B) and Chl *f* obtained with quantum chemical calculations using TDDFT with the wB-97X-V XC-functional and the pcseg-1 basis set. The difference dipole moment is shown as green arrow. The red line is the direction of the $N_B$-$N_D$ axis.

| $m$ | $\Delta E_m$ (cm$^{-1}$) | $m$ | $\Delta E_m$ (cm$^{-1}$) |
|---|---|---|---|
| $P_A$ | -240.53 | $P_B$ | - |
| $A_{-1A}$ | 71.22 | $A_{-1B}$ | 48.10 |
| $A_{0A}$ | -18.34 | $A_{0B}$ | -20.92 |

**table S6:**

Electrochromic shifts for Chl *a* in the reaction center binding sites calculated within the improved point-dipole approximation, taking into account the transition dipole moment as obtained from the quantum chemical calculations.

| $m$ | $\Delta E_m$ (cm$^{-1}$) | $m$ | $\Delta E_m$ (cm$^{-1}$) |
|---|---|---|---|
| $P_A$ | -15.34 | $P_B$ | - |
| $A_{-1A}$ | 10.86 | $A_{-1B}$ | -7.86 |
| $A_{0A}$ | 3.51 | $A_{0B}$ | 5.66 |

**table S7:**
Same as Table 6 but for Chl *f*.

Going beyond the point-dipole approximation in the case of Chl *f*, leads to a significant increase in magnitude of electrochromic shifts and in some cases to a sign switch (compare Tables S7 and S2).

Site energies of the Chl *f* pigments

| *m* | $E_m$ (nm) |
|---|---|
| $A_{-1B}$ | 747 |
| B37 | 800 |
| B38 | 800 |
| B30 | 755 |
| A23 | 759 |
| A21 | 747 |
| L03 | 738 |

**Table S8:** Site energies of the Chl *f* pigments used in the calculation of the $P^+$- P difference spectrum.

In Table S8 the site energies of the Chl *f* pigments in the antenna used in the calculation of the $P700^+$ *minus* P700 difference spectrum are given. The value for $A_{-1B}$ in the RC was obtained directly from the main S-shaped peak around 755 nm in the $P^+$- P difference spectrum. The site energies of the B37/B38 dimer are chosen such as to take into account short-range contributions resulting from coupling to charge transfer states in an effective way, as discussed above and in the main text. The site energies of B30 and B07 are taken such that the small negative dip at 662 nm in the $P700^+$ *minus* P700 difference spectrum can be reproduced. Finally, the site energies of A21, A23, and L03 were chosen such that they are in qualitative agreement with the earlier analysis of the red tail of the absorption band of FRL-PSI (*4*) Note, however, that their exact values do not matter in the present work.

In the following we apply the electrochromic shifts, determined above, to calculate the $P^+$-P difference spectrum of FR-PSI reported by Nürnberg et al., 2018.

First, we model the spectrum using the Chl *f* locations proposed in the present work, i.e. the antenna positions in table S4 and the $A_{-1B}$ position in the reaction center. The $P700^+$ *minus* P700 difference spectrum is shown in Figure 2F of the main text and the site energies of the reaction center pigments used in these calculations for PSI with neutral (P) and oxidized ($P^+$) special pair pigment $P_B$, are given in table S9.

| *m* | $E_m$ (P) (nm) | $E_m$ ($P^+$) (nm) | *m* | $E_m$ (P) (nm) | $E_m$ ($P^+$) (nm) |
|---|---|---|---|---|---|
| $P_A$ | 685.0 | 665.3 | $P_B$ | 687.0 | 720.0 |
| $A_{-1A}$ | 683.3 | 687.7 | $A_{-1B}$ | 747.0 | 749.9 |
| $A_{0A}$ | 684.5 | 684.7 | $A_{0B}$ | 685.3 | 686.6 |

**table S9:** Wavelengths corresponding to site energies of the FRL-PSI reaction center with Chl *f* in the $A_{-1B}$ binding site for PSI with neutral (P) and oxidized ($P^+$) P700 pair, with the charge localized on $P_B^+$, used in the calculation of the $P^+$- P difference spectrum in Fig. 2F of the main text. The site energy of $m = P_B$ refers to the absorption of the $P_B^+$ cation. Note, that $P_A$ also contains a site energy shift resulting from short-range contributions, as discussed in the text below.

All features of the experimental spectrum are reproduced almost quantitatively (Fig. 2F). These calculations provide additional evidence that there must be a Chl *f* in the reaction center at the $A_{-1B}$ site. In the calculations, we assume a FWHM of 150 $cm^{-1}$ for the distribution function of site energies of the reaction center pigments and the Chl *f* antenna pigments (table S4). Only for B30 a FWHM of 100 $cm^{-1}$ is assumed as suggested by the fitting of the absorption peaks done in Nürnberg et al. (*4*). In order to match the width of the linear absorption spectrum, the width of the distribution function of the Chl *a* antenna pigments is set to 250 $cm^{-1}$. The excitonic coupling within the P700 pair is increased to 225 $cm^{-1}$ as a result of the charge transfer interactions, as discussed above. These charge transfer interactions further result in an enlarged local Huang-Rhys factor of 5.2 for $P_B$ in comparison to 1.3 of $P_A$ (and the remaining RC pigments). At present this difference is just a result of the fit. Its explanation awaits a more detailed analysis of the coupling between exciton and CT states. Calculations on the charge transfer interactions in the special pair have shown (Madjet et al., 2009) that the site energies of the dimer are red-shifted by about 400 $cm^{-1}$ in comparison to that of monomeric Chl *a*. In the present calculations we assume that $P_A$ loses the CT interactions, if $P_B$ is oxidized. Consequently, its site energy is blue-shifted by about 400 $cm^{-1}$ plus the -44 $cm^{-1}$ resulting from the electrochromic shift by $P_B^+$ (table S2). The absorption band in the region between 720 nm and 740 nm is modelled by the broad absorption band of $P_B^+$ as discussed above. In fig. S20 the P700$^+$ *minus* P700 spectrum, calculated without this absorption band, is compared with the experimental spectrum. All experimental features are still reproduced, except for the broad positive band between 720 and 740 nm. In fig. S21 the difference spectrum was calculated by taking into account the absorption of P700$^+$, but increasing the inhomogeneous width fourfold from 150 $cm^{-1}$ to 600 $cm^{-1}$ in addition to the enlarged Huang-Rhys factor. In comparison to fig. S20 part of the experimental positive band between 720 nm and 740 nm is now reproduced but the agreement is less than in Fig 2F of the main text.

P700+ minus P700 difference spectrum of FR-PSI without Chl f in the reaction center

We have calculated the P700$^+$ *minus* P700 difference spectrum with the same Chl *f* positions in the antenna (table S4) but assume that there is no Chl *f* in the reaction center (fig. S13). The site energies of the reaction center pigments used in the calculations are the same as in Table S8, except for the Chl *a* at $A_{-1B}$, which has a transition wavelength of 683.3 nm and 682.5 nm in the states P700 and P700$^+$, respectively. The missing bleaching in the calculated spectrum around 750 nm demonstrates that Chl *f* at $A_{-1B}$ is responsible for this band in the experimental spectrum. The small negative peak at around 760 nm arising from the Chl *f* antenna pigments is still present.

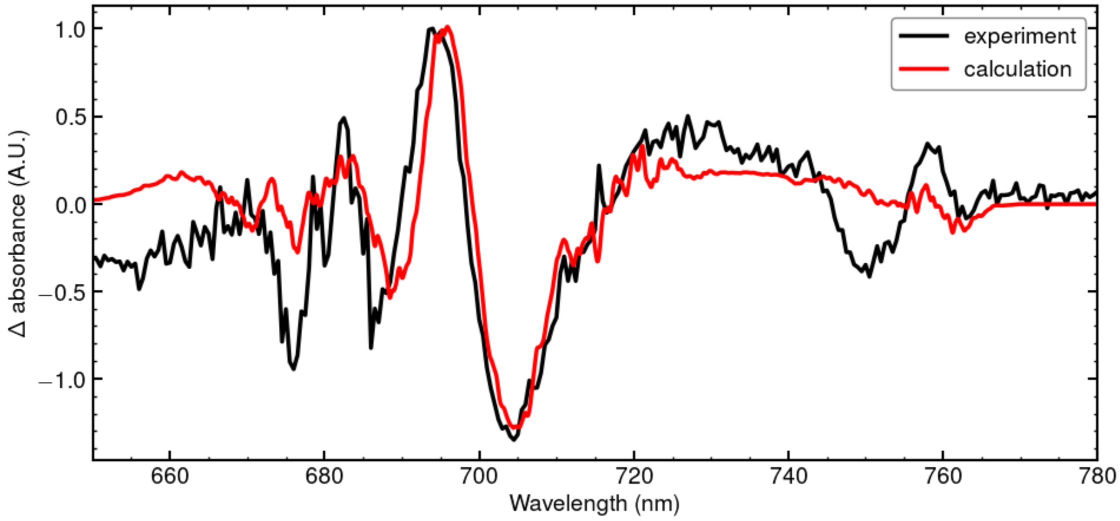


**fig. S13**

The P700$^+$ *minus* P700 difference spectrum calculated with Chl *f* located at the antenna positions based on the present cryo-EM structure, but without the Chl *f* at $A_{-1B}$ in the reaction center, is compared to experimental data.

P$^+$- P difference spectrum of FRL-PSI without Chl f in the reaction center but on B24

As a further check we calculate the P700$^+$ *minus* P700 difference spectrum without Chl *f* in the reaction center but assuming Chl *f* at the B24 binding site (assuming a site wavelength of 686 nm), the position in the antenna that gives the largest electrochromic shift. The site energies are assumed to be the same as in fig. S13.

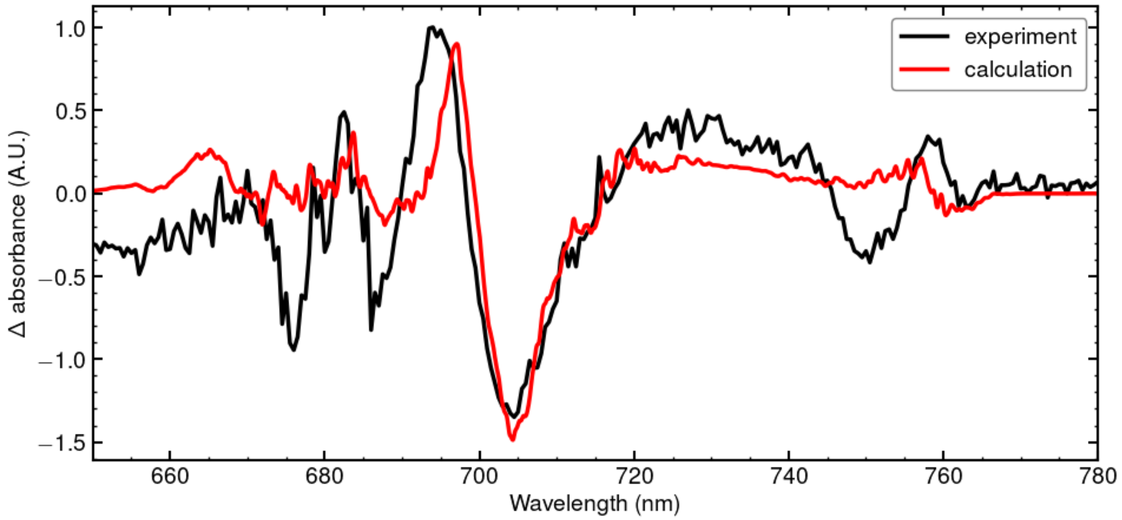


**fig. S14**

The P700$^+$ *minus* P700 difference spectrum calculated with Chl *f* located at binding sites as proposed by the present cryo-EM model, but without the Chl *f* at $A_{-1B}$, but at B24, is compared with experimental data.

Although a small peak is visible in the Chl *f* region now, the magnitude is too small to explain the experimental spectrum and the overall agreement between theory and experiment is poor. As B24 is now lacking in the Chl *a* spectral region, the fit is worse in this region too, since B24 is excitonically coupled to the RC pigments, in particular to $P_A$ and $P_B$ with a strength of 9 cm$^{-1}$ and -22 cm$^{-1}$, respectively. This coupling is effectively removed by shifting B24 out of resonance to the RC pigments. The crucial point of fig. S14 is that even the largest electrochromic shift found in the antenna is still not large enough to explain the experimental spectrum in the Chl *f* spectral region. The only possible way to obtain an electrochromic shift in the antenna that is large enough to explain the lineshape around 750 nm would be that all of the strongly shifted pigments in the antenna (B39, A40, B24) are Chl *f* and have exactly the same site energy. This assumption is not realistic and is also inconsistent with the present cryo-EM model. Therefore, the previous claims that there is no Chl *f* in the reaction center of FR-PSI can be ruled out based on the theoretical modelling of the spectroscopic data.

P700+ minus P700 difference spectrum of FR-PSI with Chl f at $A_{-1A}$

As $A_{-1A}$ is the site with the strongest electrochromic shift in the reaction center outside the special pair, we also investigate the $P700^+$ *minus* P700 difference spectrum with Chl *f* at position $A_{-1A}$ instead of $A_{-1B}$. The experimental data can be semi-quantitatively described, using the site energies in table S11, but the fit (fig. S15) is not as good as that obtained for Chl *f* at $A_{-1B}$ (fig. 2F).

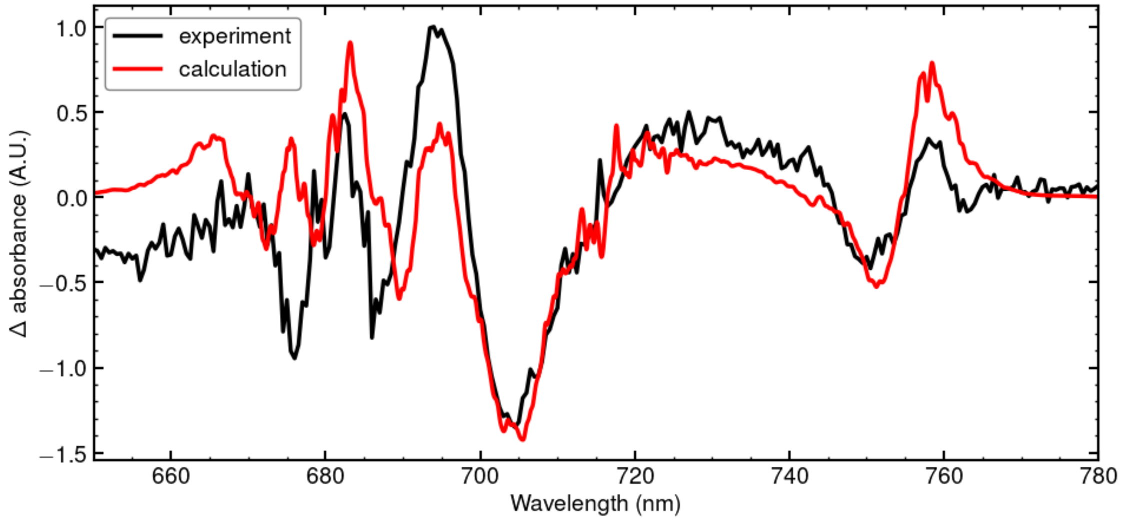


**fig. S15**
$P700^+$ *minus* P700 difference spectrum calculated with Chl *f* located at the binding sites suggested by the present cryo-EM model but with Chl *f* on $A_{-1A}$ instead of $A_{-1B}$.

| *m* | $E_m$ (P) (nm) | $E_m$ ($P^+$) (nm) | *m* | $E_m$ (P) (nm) | $E_m$ ($P^+$) (nm) |
|---|---|---|---|---|---|
| $P_A$ | 685 | 665.3 | $P_B$ | 687 | 720- |
| $A_{-1A}$ | 747 | 750.9 | $A_{-1B}$ | 683.3 | 682.5 |
| $A_{0A}$ | 684.5 | 684.7 | $A_{0B}$ | 685.3 | 686.6 |

**table S10**
Site energies fitted for the FRL-PSI reaction center pigments with Chl *f* at $A_{-1A}$.

P700+ minus P700 difference spectrum of FR-PSI at room temperature

The P700$^{+}$ *minus* P700 difference spectrum measured at room temperature showed only weak features (Nürnberg et al 2018) or no sign (*9*) of bandshifts attributable to Chl *f*. Calculations with the parameters from Table S8 are shown in fig. S12. The only difference between the parameters used in fig. S16 and fig. 2F is that at room temperature the dielectric constant ($\epsilon_{eff}$) needs to be increased to $\epsilon_{eff} = 8$ compared to the value $\epsilon_{eff}$ = 2-3 at T < 170 K (*65*). The decrease of $\epsilon_{eff}$ at low temperature is due to the freezing out of many conformational degrees of freedom. Only a very weak poorly resolved ripple is seen in the location of the 750 nm bandshift in the calculated room temperature spectrum. Note that while there are no obvious characteristic band-shift features at 750 nm despite Chl *f* being in the position of $A_{-1B}$, the other characteristic features in the experimental spectrum were qualitatively reproduced by our calculations, notwithstanding some significant offsets, the reasons of which are presently unclear. At first glance, it looks like there could be an experimental baseline drift.

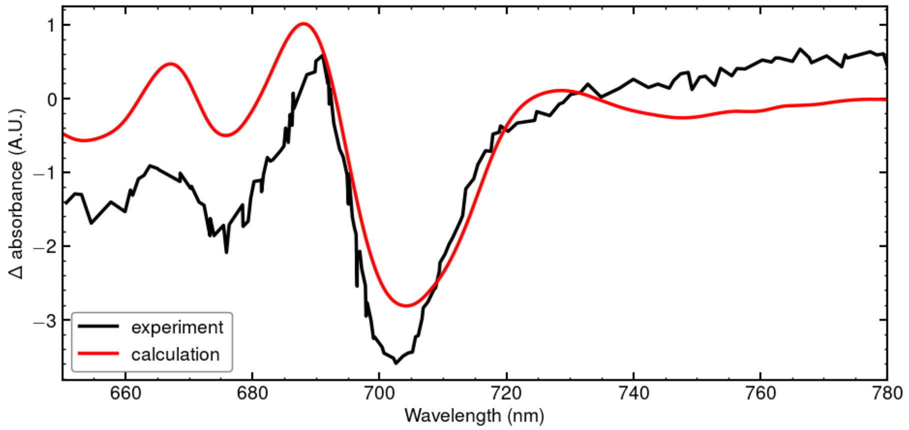


**fig. S16**
Same as in Figure 2F in the main text but at room temperature. The experimental data were taken from Kato et al. (*9*).

Additional analyses of different contributions to the P700+ minus P700 difference spectrum

Here we point out an additional important subtlety in the interpretation of the P700$^+$ *minus* P700 difference spectrum. The Chl *f* region exhibits a traditional electrochromic lineshape, as expected from a two-level system. Due to the off-resonant transition energy of the Chl *f* in the $A_{-1B}$ site with respect to the remaining RC pigments, this pigment gets effectively decoupled. This is demonstrated in fig. S17, where the difference spectrum calculated by neglecting all excitonic couplings is compared with the experimental data. While the Chl *f* region fits well, the intensities of the calculated negative peaks in the Chl *a* region are too large by a factor 2-3. Overall, it appears that the electrochromic shifts determine the band positions and the excitonic effects redistribute oscillator strength between the bands.

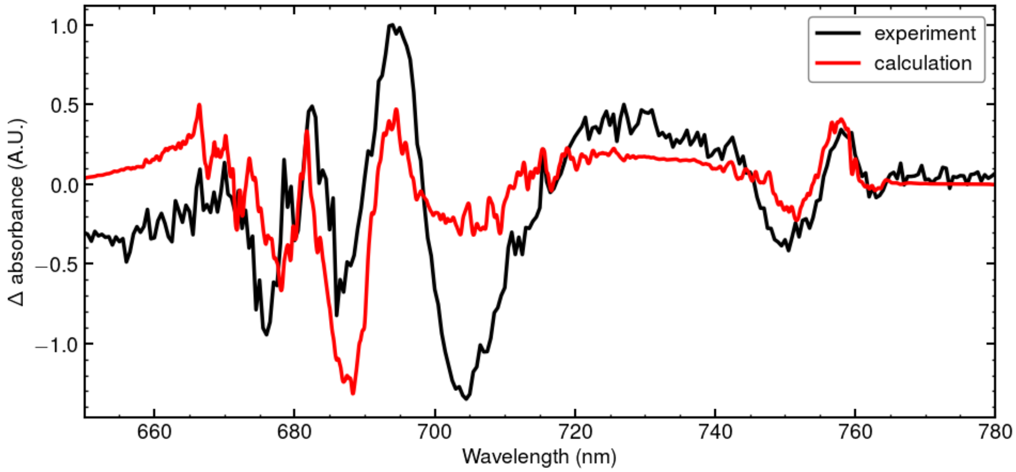


**fig. S17:**
The P700$^+$ *minus* P700 difference spectrum calculated by neglecting all excitonic couplings is compared with the experimental data.

Further evidence for this effect is obtained by a calculation in which the couplings between the pigments are included, but all electrochromic shifts are neglected (fig. S18). The only notable feature remaining in the calculated spectrum in the Chl *a* region is the bleach around 700 nm, which is due to the lower exciton state of the P700 special pair. Note that for oxidized $P_B$, the $P_A$ monomer is assumed to absorb blue-shifted at around 665 nm (table S8) due to the loss of CT couplings upon formation of $P_B^+$. The calculated spectrum also reveals a small bleaching around 755 nm, indicating that there is a small but noticeable redistribution of oscillator strength between $P_B$ and $A_{-1B}$ by their excitonic coupling (which is not present in the state $P_B^+$), despite their off-resonance.

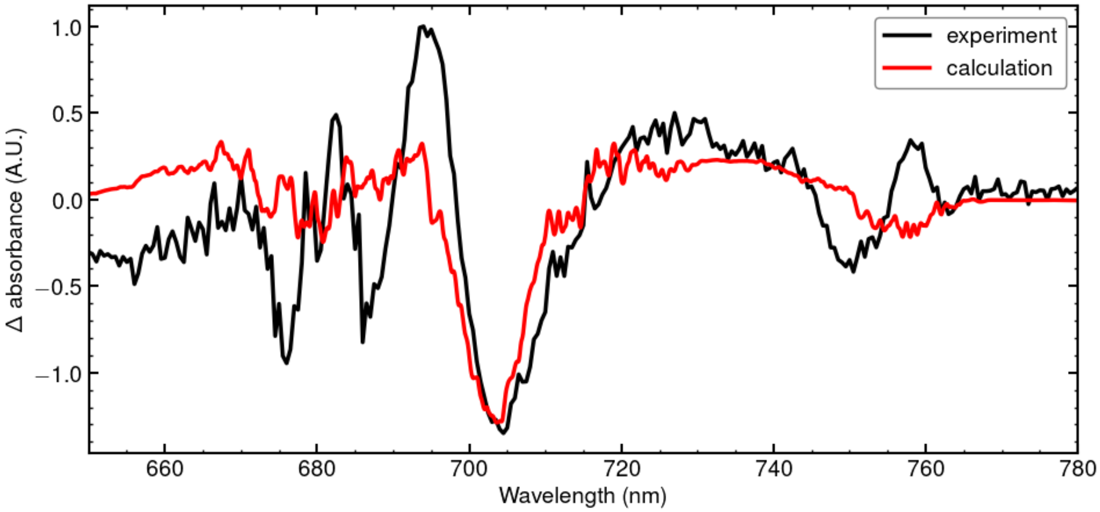


**fig. S18**
The P700$^+$ *minus* P700 difference spectrum calculated by including the excitonic couplings but neglecting all electrochromic shifts.

In order to re-establish the line positions and intensities in the Chl *a* region, we re-introduced the electrochromic shifts starting with the reaction center pigments. The resulting difference spectrum is compared in fig. S19 with the experimental data.

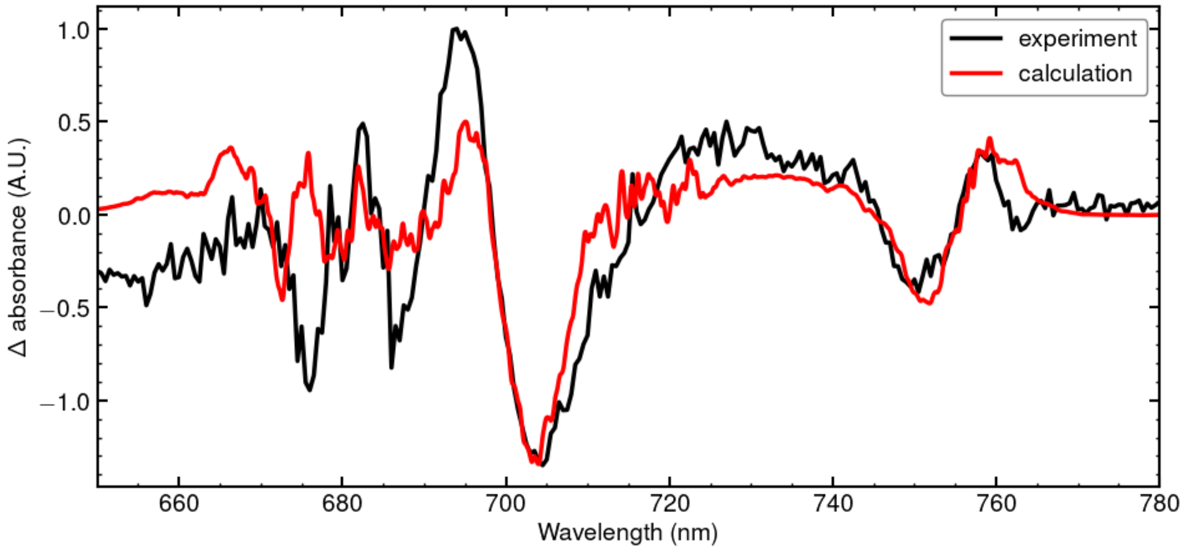


**fig. S19**

The P700$^+$ *minus* P700 difference spectrum calculated by taking into account the excitonic coupling and the electrochromic shifts of the reaction center pigments but neglecting the electrochromic shifts of the antenna pigments is compared with the experimental data.

A comparison of the calculated spectra in fig. S19 and fig. S18 shows that the electrochromic shifts of the RC pigment lead to the bands at 682 nm, 687 nm and 695 nm, but the negative band at 677 nm is still missing in the calculated spectra.

Finally, we have included the electrochromic shifts of the antenna pigments but neglected those of the RC pigments in the calculation of the difference spectrum (fig. S20). Interestingly, the whole Chl *a* region now fits qualitatively the experimental data. Hence, we have to conclude that the electrochromic shifts of the antenna pigments A40, B39 and B24 (table S1) play a significant role for the quantitative explanation of the Chl *a* region of the P700$^+$ *minus* P700 difference spectrum, outside of the main bleaching at 700 nm from the $P_AP_B$ special pair. Obviously, these antenna positions must be occupied by Chl *a*, providing further evidence that Chl *f* cannot be bound at these positions. The double bleach at 750 nm and 760 nm is a result of the electrochromic shifts of B30 and B07 and the excitonic coupling between $A_{-1B}$ and $P_B$ (seen in fig. S18).

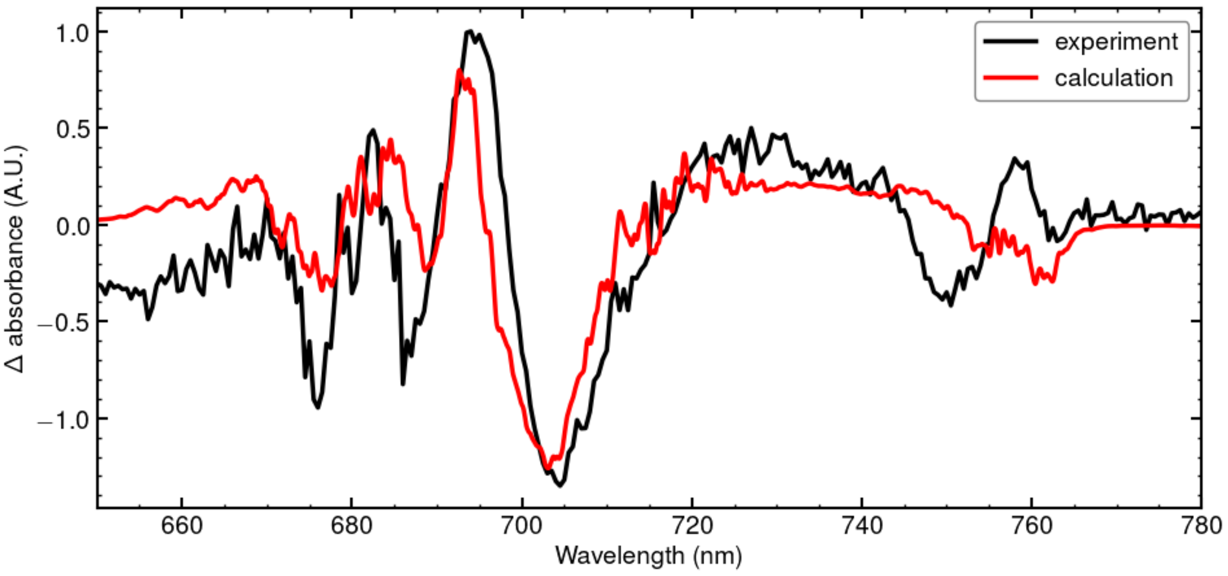


**fig. S20**

The P700$^+$ *minus* P700 difference spectrum calculated by neglecting the electrochromic shifts of the RC pigments, but taking into account the electrochomic shifts of the antenna pigments is compared with experimental data.

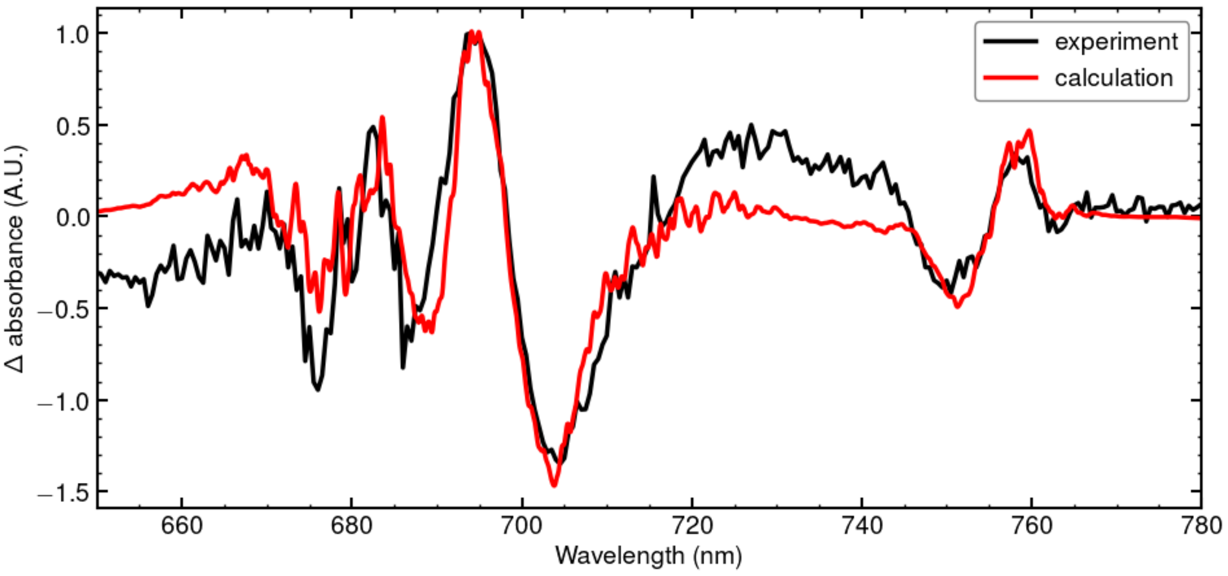


**fig. S21**

The P700$^+$ *minus* P700 difference spectrum calculated without the P700$^+$ absorption band around 730 nm is compared to experimental data.

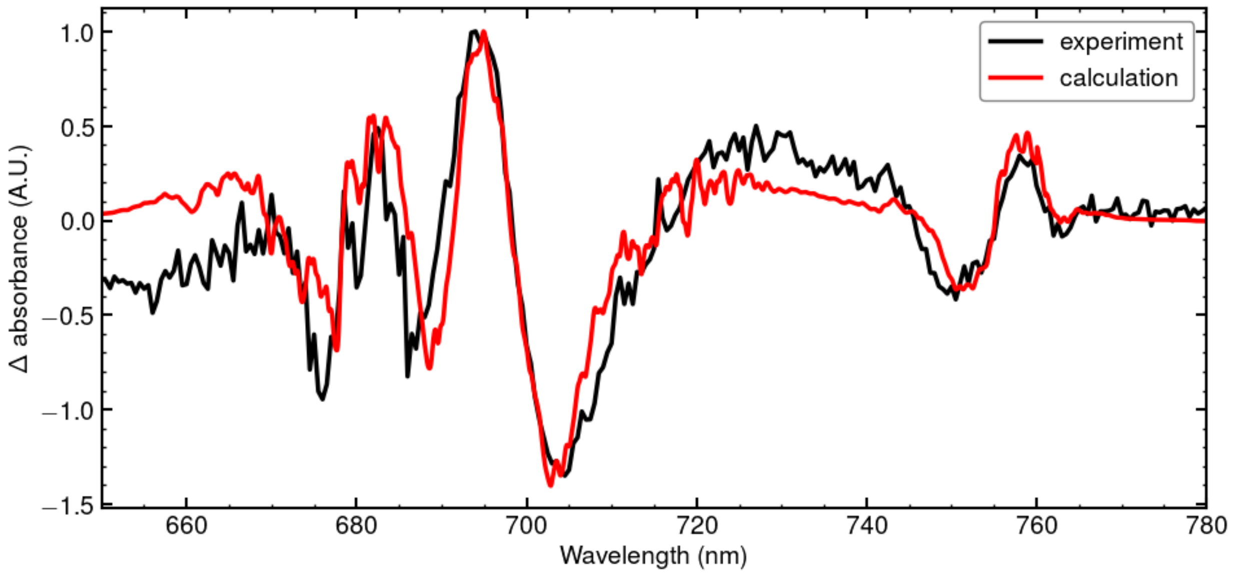


**fig. S22**

The P700⁺ *minus* P700 difference spectrum calculated with the P700⁺ absorption band around 730 nm, but with four times larger FWHM of the static disorder as in fig. 2F in the main text.

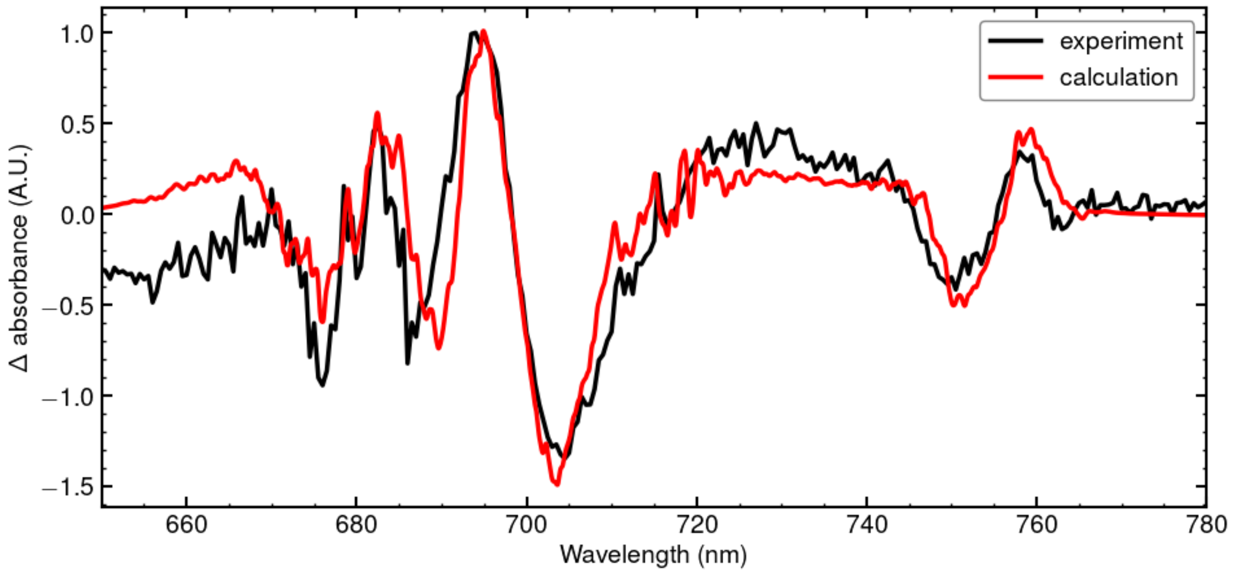


**fig. S23**

$P700^+$ *minus* P700 difference spectrum calculated interchanged site energies of B07 and B30.

In fig. S22 the $P700^+$ *minus* P700 difference spectrum is shown with interchanged site energies of B07 and B30 in comparison to the main text as both are candidates for causing the small negative peak around 760 nm referring to their electrochromic shifts. However, in this case the peak is even smaller and shifted to longer wavelengths. Shifting the site energy further to the blue to compensate for the offset results in the disappearance of the peak. These calculations further indicate that B07 is resonant with the reaction center while B30 is red-shifted.

Atomic partial charges used in the calculations of excitonic couplings and electrochromic shifts

| I | $q_I(1,0)$ | $\Delta q_I\ (e,g)$ |
|---|---|---|
| MG | 0.023748 | -0.000184 |
| CHA | -0.2166 | 0.000259 |
| CHB | 0.035896 | 0.018259 |
| CHC | 0.160303 | -0.043863 |
| CHD | -0.061769 | -0.173694 |
| NA | -0.059491 | -0.002516 |
| C1A | 0.157969 | -0.002023 |
| C2A | -0.009407 | 0.009963 |
| C3A | -0.003344 | -0.007033 |
| C4A | -0.044414 | 0.01449 |
| CMA | -0.01083 | 0.000999 |
| CAA | 0.015845 | 0.001468 |
| CBA | -0.006684 | -0.001848 |
| CGA | 0.006456 | 0.002745 |
| O1A | 3.017186 | -0.000676 |
| O2A | 0.002513 | 6.20E-05 |
| NB | 0.084869 | 0.018647 |
| C1B | -0.09321 | -0.036598 |
| C2B | 0.015615 | -0.024023 |
| C3B | 0.02176 | -0.025548 |
| C4B | -0.157577 | 0.033194 |
| CMB | -0.02764 | 0.001712 |
| CAB | -0.014147 | 0.000859 |
| CBB | -0.020941 | -0.005628 |
| NC | 0.051317 | -0.069774 |
| C1C | -0.003699 | 0.03968 |
| C2C | 0.024722 | 0.018701 |
| C3C | -0.003838 | -0.021956 |
| C4C | 0.337839 | 0.17353 |
| CMC | -0.000535 | 0.00464 |
| CAC | -0.012029 | -0.001643 |
| CBC | -0.003703 | 0.000837 |
| ND | -0.16459 | -0.04216 |
| C1D | 0.128725 | 0.137883 |
| C2D | 0.449065 | -0.044596 |
| C3D | 0.22094 | -0.008733 |
| C4D | 0.25355 | 0.04683 |
| CMD | 0.039975 | -0.000355 |
| CAD | 0.139862 | -0.003223 |

| | | |
|---|---|---|
| OBD | 0.014565 | -0.002589 |
| CBD | 0.017518 | -0.006256 |
| CGD | -0.012368 | 0.003348 |
| O1D | 0.012282 | -0.000856 |
| O2D | 0.000684 | -0.001407 |
| CED | 0.005975 | -7.60E-05 |
| C1 | -0.000241 | -0.000846 |

Table S11: Transition and difference charges of Chl *a* in units of the elementary charge $\boldsymbol{e}$.

| I | $\boldsymbol{q_I(1,0)}$ | $\boldsymbol{\Delta q_I\ (e,g)}$ |
|---|---|---|
| MG | 0.02422 | -0.001302 |
| CHA | -0.19909 | 0.024801 |
| CHB | 0.05913 | -0.002097 |
| CHC | 0.13745 | -0.12499 |
| CHD | -0.10386 | -0.13245 |
| NA | -0.0445 | -0.00322 |
| C1A | 0.15035 | -0.00934 |
| C2A | -0.00975 | 0.005269 |
| C3A | -0.00606 | -0.001 |
| C4A | -0.04587 | 0.027586 |
| CMA | -0.00701 | 0.001309 |
| CAA | 0.01902 | 0.002159 |
| CBA | -0.00635 | -0.00186 |
| CGA | 0.00617 | 0.003609 |
| O1A | -0.00562 | -0.00127 |
| O2A | 0.00206 | -0.00016 |
| NB | 0.08312 | -0.01042 |
| C1B | -0.10649 | 0.019879 |
| C2B | 0.02713 | -0.03502 |
| C3B | 0.01404 | -0.0369 |
| C4B | -0.1493 | 0.091375 |
| C467 | -0.03834 | -0.00063 |
| O370 | -0.00403 | -0.00423 |
| CAB | -0.02072 | 0.000982 |
| CBB | -0.04866 | -0.00734 |
| NC | 0.02469 | -0.08279 |
| C1C | -0.13014 | 0.125103 |
| C2C | 0.02159 | 0.005418 |
| C3C | -0.01198 | -0.00245 |
| C4C | 0.08857 | 0.137846 |
| CMC | 0.00981 | 0.003062 |

| | | |
|---|---|---|
| CAC | -0.00659 | 0.000725 |
| CBC | -0.0015 | 0.001162 |
| ND | -0.1699 | -0.01073 |
| C1D | 0.14363 | 0.096526 |
| C2D | -0.00283 | -0.04565 |
| C3D | -0.05025 | -0.01554 |
| C4D | 0.2509 | -0.00136 |
| CMD | 0.03533 | -0.00127 |
| CAD | 0.03801 | -0.00102 |
| OBD | 0.01356 | -0.00444 |
| CBD | 0.01144 | -0.00882 |
| CGD | -0.01271 | 0.002206 |
| O1D | 0.01206 | -0.00021 |
| O2D | 0.00118 | -1.55E-03 |
| CED | 0.00711 | -3.90E-05 |
| C1 | 0.00097 | -0.000929 |

Table S12: Transition and difference charges of Chl *f* in units of the elementary charge $\boldsymbol{e}$.

| I | $\boldsymbol{q_I(1,0)}$ | $\boldsymbol{\Delta q_I\ (g_+, g)}$ |
|---|---|---|
| MG | - | 1.042065 |
| CHA | - | 1.041999 |
| CHB | - | -1.243017 |
| CHC | - | -0.282684 |
| CHD | - | -0.722535 |
| NA | - | -1.759463 |
| C1A | - | 0.412689 |
| C2A | - | -0.391993 |
| C3A | - | 0.167983 |
| C4A | - | 1.94441 |
| CMA | - | -0.046369 |
| CAA | - | -0.046581 |
| CBA | - | -0.000267 |
| CGA | - | -0.684972 |
| O1A | - | 0.501469 |
| O2A | - | 0.492436 |
| NB | - | -1.110973 |
| C1B | - | 1.046962 |
| C2B | - | 0.031079 |
| C3B | - | -0.46416 |
| C4B | - | 0.852368 |
| CMB | - | 0.090619 |

| | | |
|---|---|---|
| CAB | - | 0.114099 |
| CBB | - | 0.037454 |
| NC | - | -0.652058 |
| C1C | - | 0.381364 |
| C2C | - | -0.08708 |
| C3C | - | -0.148624 |
| C4C | - | 0.860398 |
| CMC | - | 0.086575 |
| CAC | - | 0.032201 |
| CBC | - | 0.064949 |
| ND | - | -1.083271 |
| C1D | - | 1.10731 |
| C2D | - | 0.139026 |
| C3D | - | -0.811267 |
| C4D | - | 0.45274 |
| CMD | - | 0.039896 |
| CAD | - | 0.608529 |
| OBD | - | -0.003851 |
| CBD | - | -0.710177 |
| CGD | - | -0.555956 |
| O1D | - | 0.4966 |
| O2D | - | 0.36829 |
| CED | - | -0.293832 |
| C1 | - | -0.314376 |

Table S13: Difference charges of $Chl^{+}$ $a$ in units of the elementary charge $\boldsymbol{e}$. No transition charges are given as the $P_B^{+}$ is assumed to be excitonically fully decoupled because of the small dipole strengths and the red-shifted absorption of the cation.

| I | $\boldsymbol{q_I(1,0)}$ | $\boldsymbol{\Delta q_I\ (e,g)}$ |
|---|---|---|
| MG | 0.023897 | -0.00265 |
| CHA | -0.217318 | -0.006749 |
| CHB | 0.035915 | 0.003621 |
| CHC | 0.16029 | -0.054391 |
| CHD | -0.0618 | -0.153248 |
| NA | -0.059759 | -0.011859 |
| C1A | 0.158694 | 0.010089 |
| C2A | -0.009593 | 0.010415 |
| C3A | -0.003478 | -0.00421 |
| C4A | -0.044453 | 0.019757 |
| CMA | -0.010772 | 0.001395 |
| CAA | 0.01585 | 0.001903 |

| | | |
|---|---|---|
| CBA | -0.006232 | -0.001548 |
| CGA | 0.006371 | 0.003623 |
| O1A | -0.005863 | -0.001364 |
| O2A | 0.002128 | -0.001156 |
| NB | 0.084792 | 0.016348 |
| C1B | -0.09317 | -0.025989 |
| C2B | 0.015623 | -0.029386 |
| C3B | 0.021746 | -0.025026 |
| C4B | -0.157531 | 0.035216 |
| CMB | -0.027644 | 0.001585 |
| CAB | -0.014148 | 0.000319 |
| CBB | -0.020942 | -0.007032 |
| NC | 0.051258 | -0.067526 |
| C1C | -0.163313 | 0.051598 |
| C2C | 0.024753 | 0.017754 |
| C3C | -0.003885 | -0.017569 |
| C4C | 0.03235 | 0.160219 |
| CMC | -0.000541 | 0.004741 |
| CAC | -0.012019 | -0.001562 |
| CBC | -0.003708 | 0.000846 |
| ND | -0.164841 | -0.033922 |
| C1D | 0.128821 | 0.124115 |
| C2D | -0.00817 | -0.043641 |
| C3D | -0.034514 | -0.006307 |
| C4D | 0.253956 | 0.049326 |
| CMD | 0.039986 | -0.000331 |
| CAD | 0.028543 | -0.010752 |
| OBD | 0.014548 | -0.003824 |
| CBD | 0.017547 | -0.003235 |
| CGD | -0.012293 | 0.002641 |
| O1D | 0.012273 | -0.000893 |
| O2D | 0.000681 | -0.001166 |
| CED | 0.005962 | -0.000176 |

Table S14: Transition and difference charges for the Chl *a* without the C1 atom in units of the elementary charge $\boldsymbol{e}$.

| I | $\boldsymbol{q_I(1,0)}$ | $\boldsymbol{\Delta q_I\,(e,g)}$ |
|---|---|---|
| MG | 0.024652 | -0.001666 |
| CHA | -0.19998 | 0.021237 |
| CHB | 0.067771 | 0.009236 |
| CHC | 0.134197 | -0.121236 |

| | | |
|---|---|---|
| CHD | -0.10442 | -0.132968 |
| NA | -0.03921 | 0.004816 |
| C1A | 0.149564 | -0.008287 |
| C2A | -0.00931 | 0.005551 |
| C3A | -0.00512 | 0.000412 |
| C4A | -0.05493 | 0.013168 |
| CMA | -0.00703 | 0.001533 |
| CAA | 0.018927 | 0.001932 |
| CBA | -0.00634 | -0.001366 |
| CGA | 0.00629 | 0.003186 |
| O1A | -0.00565 | -0.001088 |
| O2A | 0.0021 | -0.000184 |
| NB | 0.081412 | -0.011596 |
| C1B | -0.1137 | 0.01348 |
| C2B | 0.038526 | -0.027135 |
| C3B | 0.004044 | -0.038484 |
| C4B | -0.14204 | 0.092066 |
| C467 | -0.04681 | -0.008995 |
| CAB | -0.0162 | -0.001346 |
| CBB | -0.05079 | -0.006777 |
| NC | 0.023386 | -0.079711 |
| C1C | -0.12889 | 0.119199 |
| C2C | 0.022344 | 0.007167 |
| C3C | -0.01315 | -0.002587 |
| C4C | 0.08979 | 0.137755 |
| CMC | 0.009725 | 0.002758 |
| CAC | -0.00635 | 0.000718 |
| CBC | -0.00154 | 0.001113 |
| ND | -0.17148 | -0.014131 |
| C1D | 0.145192 | 0.098492 |
| C2D | -0.00399 | -0.04698 |
| C3D | -0.04902 | -0.01415 |
| C4D | 0.250935 | 0.000539 |
| CMD | 0.035483 | -0.001096 |
| CAD | 0.037413 | -0.001833 |
| OBD | 0.01366 | -0.00434 |
| CBD | 0.011967 | -0.007959 |
| CGD | -0.01298 | 0.00234 |
| O1D | 0.012272 | -0.000306 |
| O2D | 0.001249 | -0.001531 |
| CED | 0.007077 | 3.199999E-05 |
| C1 | 0.000939 | -0.000914 |

Table S15: Transition and difference charges of Chl *f*, obtained by excluding the oxygen atom that defines the difference to Chl *a* in the fit of the ab-initio ESP*,* in units of the elementary charge ***e***. These charges are used to "transform" a Chl *a* in the structural model to a Chl *f.*

Exciton domains

| Domain | Pigments |
|---|---|
| 1 | 1, 2, 3, 4, 5, 6, 7, 8, 9, 10, 14, 15, 17, 18, 19, 20, 22, 23, 24, 25, 26, 27, 28, 29, 30, 31, 32, 33, 34, 35, 36, 37, 38, 39, 40, 44, 45, 46, 47, 53, 57, 59, 60, 61, 68, 69, 73, 74, 75, 76, 77, 78, 79, 80, 81, 82, 83, 84, 85, 86, 89 |
| 2 | 11, 13, 21 |
| 3 | 12 |
| 4 | 16 |
| 5 | 41, 42, 43 |
| 6 | 48, 49, 50, 56, 72 |
| 7 | 51, 52, 70, 71 |
| 8 | 54, 55, 63 |
| 9 | 58 |
| 10 | 62, 64, 65, 66, 67 |
| 11 | 87 |
| 12 | 88 |

Table S17: Exciton domains obtained with a cut-off excitonic coupling of 30 $cm^{-1}$..

**Supplementary Text S4: $A_{-1B}$ /$A_{0B}$ Electronic Coupling**

The structural model for the $A_{-1B}$ and $A_{0B}$ chlorophylls in FR-PSI and WL-PSI show a small rotation of the chlorophylls relative to each other (*28*). Here the electronic and excitonic couplings were calculated using three different structural models for the $A_{-1B}$ /$A_{0B}$ pair: 1) Chl *a* Chl *a* using the coordinates of WL-PS1 from *H. hongdechloris* (6KMW), 2) Chl *f* Chl *a* using the coordinates from the present work) as a control Chl *a* Chl *a* using the coordinates from the present work. table S18 shows the calculated electron ($J_e$), hole ($J_h$) and exciton ($J_{ex}$) couplings between the chlorophylls of the different structures. The change in the overlap between the chlorophylls leads to larger transfer integrals between them, resulting in increased electronic coupling, which is even more significant in the case of the Chl *f* Chl *a* pair. The electronic couplings were calculated from the transfer integrals between their LUMO orbitals for the electron and HOMO orbitals for the hole, calculated using the counterpoise method (*66*). This is based on the molecular orbitals of the pair and of each chlorophyll separately, where ghost atoms have been used to ensure the same basis set. The excitonic couplings were calculated from the DFT description of the wavefunction using the electronic energy transfer (EET) analysis method in Gaussian 16. For the calculations, chlorophyll geometries were simplified by reducing chains longer than a methyl group, including the phytol tail, to a single methyl group. We considered amino acids and water molecules within 5 angstroms of the central magnesium, i.e., a water molecule and asparagine in the case of $A_{-1B}$, and methionine for $A_{0B}$, see fig. S24. The geometries were individually optimised in vacuum. All quantum chemistry calculations were performed with DFT using B3LYP/6-31g* level of theory and carried out in Gaussian 16. Transfer integrals were calculated using a home-built software.

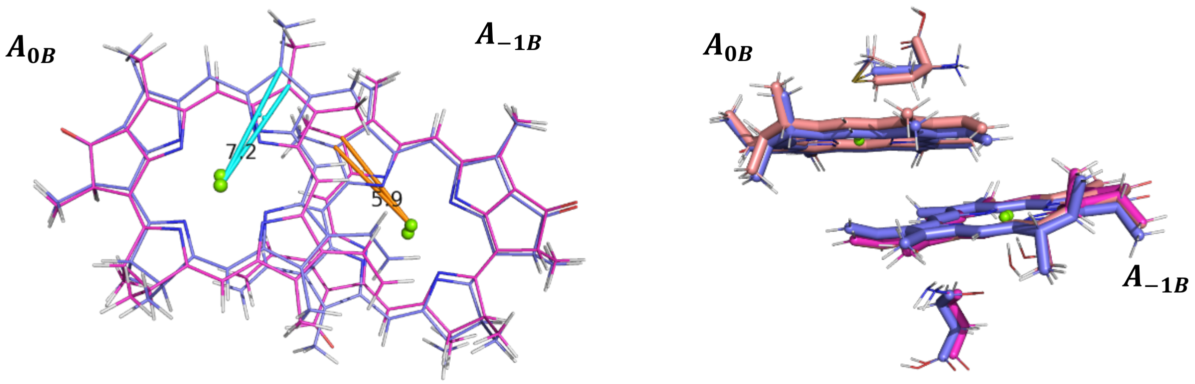


**fig. S24**

Top and front views of the overlay of the $A_{-1B}$, $A_{0B}$ dimers in the present work and the WL-PSI previously reported of *H. hongdechloris* (*9*). The top view includes auxiliary lines in cyan and orange, along with labels indicating the rotation (in degrees) of chlorophylls in this work compared to Kato et al (*9*).

| | $J_e\ (meV)$ | $J_h\ (meV)$ | $J_{ex}\ (meV)$ |
|---|---|---|---|
| **Kato (Chl *a*, Chl *a*)** | 8.9 | 10.8 | 10.7 |
| **Consoli (Chl *a*, Chl *a*)** | 17.6 | 36.6 | 8.3 |
| **Consoli (Chl *a*, Chl *f*)** | 19.8 | 66.0 | 7.8 |

**table S18**.
Calculated electron, hole and excitonic couplings between the $A_{-1B}$ and $A_{0B}$ chlorophylls. $J_e$, Jh and $J_{ex}$ are the electronic, hole and excitonic coupling values. Where both are Chl *a,* in WL-PSI, the geometry from Kato et al (2020) is used; where both are Chl *a* in FR-PSI, and when $A_{-1B}$ is Chl-f and $A_{0B}$ is Chl *a*, the geometry from the present work is used.

.

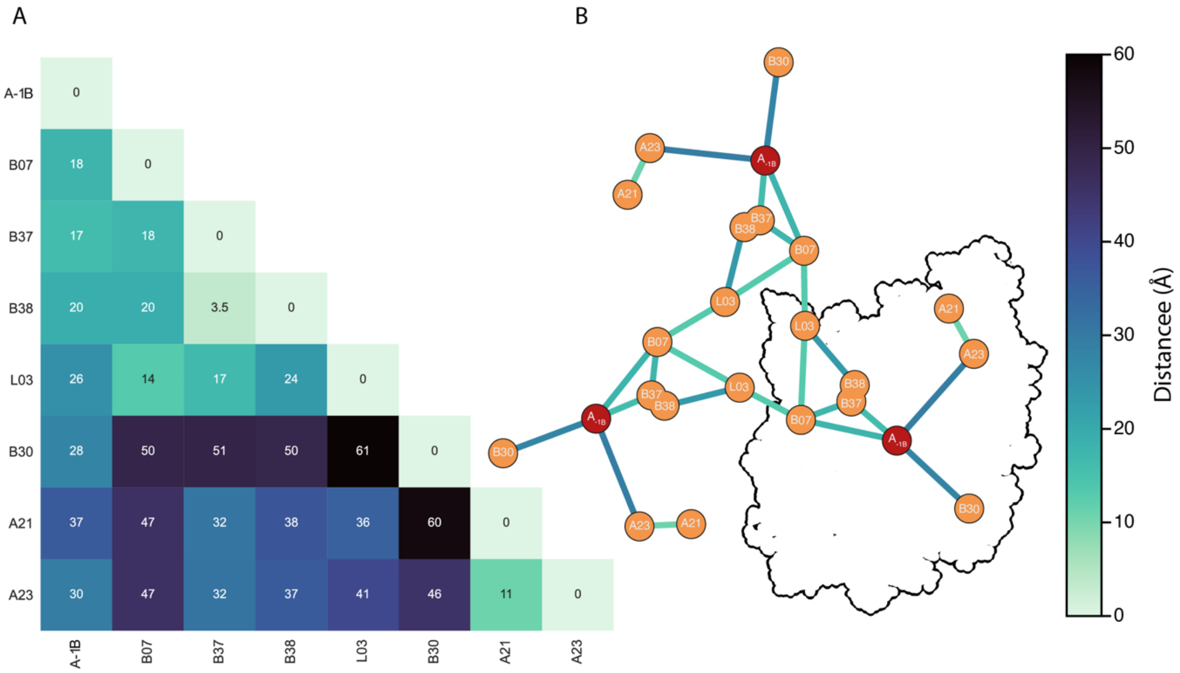


**Fig. S25**

A) Chl *f* edge-to-edge distance matrix color coded according to the colorbar on the right. B) shortest distances among Chl *f* sites showing the potential exciton routes towards the reaction centre and between the monomers in FR-PSI trimer color coded according to the colorbar on the right. FR-PSI monomer outline is shown in black, antenna Chl *f* are in orange and reaction centre $A_{-1B}$ site in red.

| Jordan et al. ID | Kato et al. ID | 7S3D | 6KMX | 9EYS | Ligand |
|---|---|---|---|---|---|
| A1011 | A801 | | | | delta |
| A1012 | A802 | | | Y | H2O |
| A1013 | A803 | | | | Met |
| A1101 | A804 | | | | delta |
| A1102 | A805 | | | | delta |
| A1103 | A806 | | | | delta |
| A1104 | A807 | | | | delta |
| A1105 | A808 | | | | delta |
| A1106 | A809 | | | | Gln114 |
| A1107 | A810 | | | | Gln122 |
| A1108 | A811 | | | | delta |
| A1109 | A812 | | | | delta |
| A1110 | A813 | | | | delta |
| A1111 | A814 | | | | delta |
| A1112 | A815 | | | | epsilon |
| A1113 | A816 | | | | delta |
| A1114 | A817 | | | | H2O |
| A1115 | A818 | | | | delta |
| A1116 | A819 | | | | epsilon |
| A1117 | A820 | | | | delta |
| A1118 | A821 | | | | delta |
| A1119 | A822 | | | | H20 |
| A1120 | A823 | | | | delta |
| A1121 | A824 | Y | | Y | H2O |
| A1122 | A825 | | | | delta |
| A1123 | A826 | SS | Y | SS | H2O |
| A1124 | A827 | | Y | | H2O |
| A1125 | A828 | | | | delta |
| A1126 | A829 | | | | epsilon |
| A1127 | A830 | | Y | | delta |
| A1128 | A831 | | | | delta |
| A1129 | A832 | | Y | | delta |
| A1130 | A833 | | | | delta |
| A1131 | A834 | | | | delta |
| A1132 | A835 | | | | epsilon |
| A1133 | A836 | | | | delta |
| A1134 | A837 | | | | Thr528 |
| A1135 | A838 | | | | delta |

| | | | | | |
|---|---|---|---|---|---|
| A1136 | A839 | | | | delta |
| A1137 | A840 | | | | delta |
| A1138 | A841 | | | | delta |
| A1139 | A842 | | | | H2O |
| A1140 | A843 | | | | delta |
| B1021 | B801 | | | | delta |
| B1022 | B802 | | | | H2O |
| B1023 | B803 | | | | delta |
| B1201 | B804 | | | | delta |
| B1202 | B805 | | | | delta |
| B1203 | B806 | | | | delta |
| | B807 | | | | delta |
| B1205 | B808 | | | | delta |
| B1206 | B809 | | | | Asp93 |
| B1207 | L202 | Y | Y | Y | Gln95 |
| B1208 | B810 | | | | delta |
| B1209 | B811 | | | | delta |
| B1210 | B812 | | | | delta |
| B1211 | B813 | | | | epsilon |
| B1212 | B814 | | | | delta |
| B1213 | B815 | | | | delta |
| B1214 | B816 | | | | epsilon |
| B1215 | B817 | | | | delta |
| B1216 | B818 | | | | H2O |
| B1217 | B819 | | | | delta |
| B1218 | B820 | | | | delta |
| B1219 | B821 | | | | H2O |
| B1220 | B822 | | | | delta |
| B1221 | B823 | | | | H2O |
| B1222 | B824 | | Y | | H2O |
| B1223 | B825 | | | | delta |
| B1224 | B826 | | | | epsilon |
| B1225 | B827 | | | | delta |
| B1226 | B828 | | | | delta |
| B1227 | B829 | | | | delta |
| B1228 | B830 | | | | delta |
| B1229 | B831 | | | | delta |
| B1230 | B832 | Y | | Y | epsilon |
| B1231 | B833 | | | | delta |
| B1232 | B834 | | | | H2O |
| B1233 | B835 | | | | H2O |

| | | | | | |
|---|---|---|---|---|---|
| B1234 | B836 | | | | delta |
| B1235 | B837 | | | | delta |
| B1236 | B838 | | | | delta |
| B1237 | A844 | Y | Y | Y | H2O |
| B1238 | B839 | Y | | Y | H2O |
| B1239 | B840 | | | | delta |
| K104 | K101 | | | | delta |
| L1501 | L204 | | | | Glu66 |
| L1502 | L205 | | | | His |
| L1503 | L206 | | | Y | H2O |
| X1701 | NON PRESENT | | | | |

**Table S18**

Chl f candidates in FR-PSI structures available together with a conversion table between the two naming conventions used for chlorophyll sites, cells with a green Y correspond to an assignement of a Chl *f* site, SS in yellow corresponds to a species-specific assignment. The Ligand column refers to the ligand to the central $Mg^{2+}$ coordination, H2O when it's coordinated by a water, aminoacid and number when they are coordinated by a non histidine aminoacid, and delta and epsilon to highlight which nitrogen in the histidine sidechain coordinates the $Mg^{2+}$.

| | | | | |
|---|---|---|---|---|
| All-Atom Contacts | Clashscore, all atoms: | 10.61 | | $68^{th}$ percentile$^{*}$ (N=1784, all resolutions) |
| | Clashscore is the number of serious steric overlaps (> 0.4 Å) per 1000 atoms. | | | |
| Protein Geometry | Poor rotamers | 0 | 0.00% | Goal: <0.3% |
| | Favored rotamers | 5603 | 99.77% | Goal: >98% |
| | Ramachandran outliers | 0 | 0.00% | Goal: <0.05% |
| | Ramachandran favored | 6716 | 98.19% | Goal: >98% |
| | Rama distribution Z-score | 1.21 ± 0.10 | | Goal: abs(Z score) < 2 |
| | MolProbity score$^{\wedge}$ | 1.54 | | $94^{th}$ percentile$^{*}$ (N=27675, 0Å - 99Å) |
| | Cβ deviations >0.25Å | 0 | 0.00% | Goal: 0 |
| | Bad bonds: | 0 / 77625 | 0.00% | Goal: 0% |
| | Bad angles: | 0 / 109056 | 0.00% | Goal: <0.1% |
| Peptide Omegas | Cis Prolines: | 9 / 351 | 2.56% | Expected: ≤1 per chain, or ≤5% |
| | Cis nonProlines: | 3 / 6531 | 0.05% | Goal: <0.05% |
| Low-resolution Criteria | CaBLAM outliers | 61 | 0.9% | Goal: <1.0% |
| | CA Geometry outliers | 29 | 0.43% | Goal: <0.5% |
| Additional validations | Chiral volume outliers | 0/9651 | | |
| | Waters with clashes | 38/399 | 9.52% | See UnDowser table for details |

**Table S19**
Results of the validation output from MolProbity
